# Automated Quantification of Stereotypical Motor Movements in Autism Using Persistent Homology

**DOI:** 10.1101/2025.09.03.674008

**Authors:** Austin A. MBaye, Jose A. Perea, Christopher J. Tralie, Matthew S. Goodwin

**Affiliations:** Department of Mathematics, Northeastern University, Boston, MA, USA; Department of Mathematics and Khoury College of Computer Sciences, Northeastern University, Boston, MA, USA; Department of Mathematics and Computer Sciences, Ursinus College, Collegeville, PA, USA; Bouve College of Health Sciences & Khoury College of Computer Sciences, Northeastern University, Boston, MA, USA

## Abstract

Stereotypical motor movements (SMM) are a core diagnostic feature of autism that remain difficult to quantify efficiently and validly across individuals and developmental stages. The current paper presents a novel pipeline that leverages Topological Data Analysis to quantify and characterize recurrent movement patterns. Specifically, we use persistent homology to construct low-dimensional, interpretable feature vectors that capture geometric properties associated with autistic SMM by extracting periodic structure from time series derived from pose estimation landmarks in video data and accelerometer signals from wearable sensors. We demonstrate that these features, combined with simple classifiers, enable accurate automated quantification of autistic SMM. Visualization of the learned feature space reveals that extracted features generalize across individuals and are not dominated by person-specific SMM. Our results highlight the potential of using mathematically principled features to support more scalable, interpretable, and person-agnostic characterization of autistic SMM in naturalistic settings.

## Introduction

Autism is a complex neurodevelopmental condition characterized by common behavioral and social characteristics that complicate social integration and impact daily living. The Diagnostic and Statistical Manual of Mental Disorders, Fifth Edition (DSM-5)[8] identifies two main diagnostic criteria for autism: (a) persistent deficits in social communication and social interaction across contexts, and (b) restricted, repetitive patterns of behavior, interests, or activities. These behavioral patterns often include stereotypical motor movements (SMM), with the most frequent forms observed in autism being hand flapping, body rocking, and finger flicking [18].

Traditional methods of measuring SMM in autism include general rating scales [23, 20], live observation, and manually annotating videos. While helpful for documenting the presence or absence of SMM, these measures rely on clinician interviews, limited behavioral observation, and parental report, all of which can be subjective, inaccurate, and difficult to compare across individuals with autism and over time. Video-based observation is more accurate than interviews, rating scales, and direct observation. However, they are time-consuming and require accurate offline annotation of videos. Moreover, both live and video measures of SMM are typically based on observations in controlled environments (i.e., a lab or clinic), rather than in natural environments (i.e., a home or classroom). More objective and scalable methods for monitoring SMM in naturalistic settings over time are needed to determine whether topography, frequency, duration, intensity, or temporal patterning changes have occurred. Accurate measures of SMM are necessary to assess their functional significance and guide appropriate interventions when needed.

In recent years, accelerometry-based systems show promise in capturing movement dynamics associated with SMM [1, 30, 33, 12]. However, they, too, have limitations. Machine learning models must contend with high intra- and inter-individual variability in the frequency, intensity, and form of SMMs, making consistent classification difficult. There is also a trade-off between achieving high accuracy and maintaining model interpretability, which is critical for clinical utility. Finally, accelerometer-based methods rely on wearable sensors, which can be uncomfortable for some individuals, particularly those with sensory sensitivities, and costly to sustain.

Methods for quantifying SMM using video would provide a noninvasive, scalable solution, suitable for long-term, real-world deployment. Although prior video-based methods show promise, most datasets [36, 19, 27, 15] are limited by their omission of clinical and demographic metadata, such as formal autism diagnoses, age, and developmental context. Furthermore, models that require complex data pre-processing to achieve high classification accuracy often have low interpretability, restricting their use in clinical environments.

To address the above challenges, we propose a novel unifying pipeline grounded in Topological Data Analysis (TDA) – AQSM - *S*W1*PerS*: Automated Quantification of Stereotypical Motor Movements via Sliding Windows and 1-Persistence Scoring – and demonstrate its utility in previously published accelerometer data and naturalistic videos from [11], which include clinical and demographic information and multiple environmental recordings. Specifically, we leverage the persistent homology of sliding window embeddings to capture autistic SMM’s intrinsic structure and periodicity directly from time series data. As illustrated in Figure 1, TDA offers a mathematically principled framework for summarizing the recurrent behavior of complex time series that is robust to noise, sampling artifacts, and individual variability. Moreover, the methodology produces interpretable, person-agnostic features with minimal preprocessing across different data modalities.

**Figure 1:**
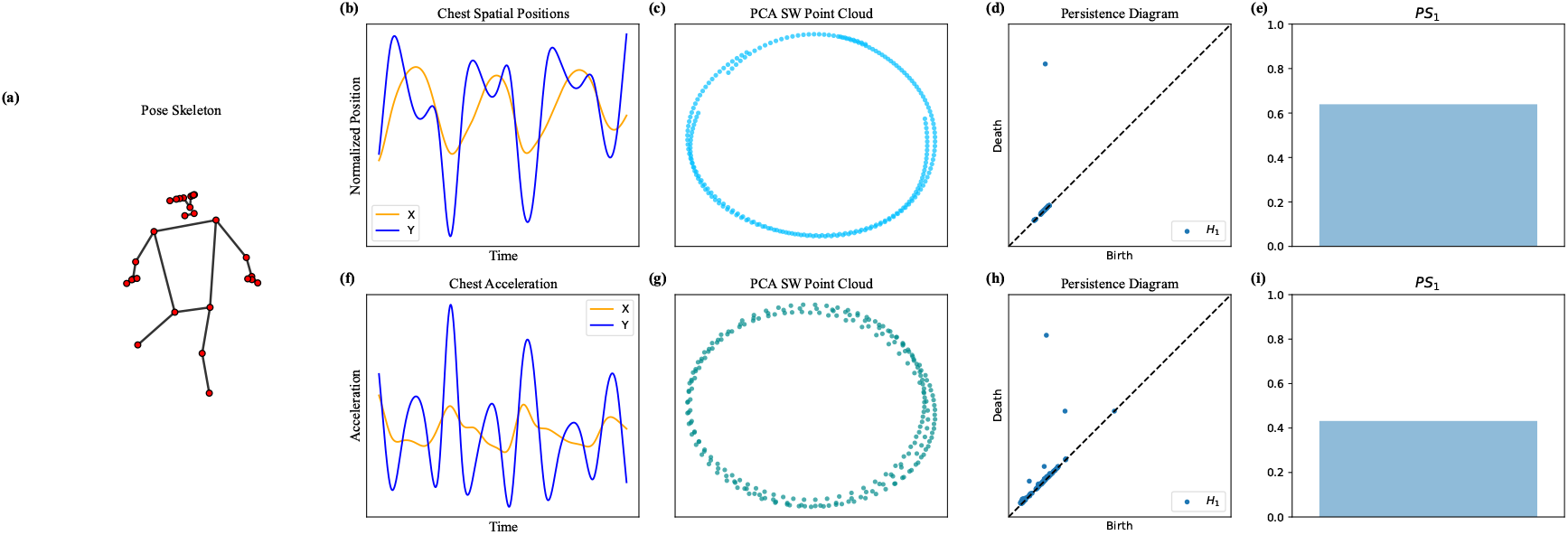
Summary of the AQSM - *S*W1*PerS* pipeline for video data. (a) fitted skeleton on an autistic participant engaged in stereotypical rocking (this step uses MediaPipe+YOLOv5), (b) chest trajectory inferred from the moving fitted skeleton, (f) acceleration representation of chest trajectory, (c, g) sliding window point clouds of respective trajectories – the periodicity of the movement corresponds to the circularity of the point cloud, (d, h) corresponding persistence diagrams measuring the circularity of the sliding window point clouds – circularity is strongest when only one point in the diagram is far up and to the left (e, i) periodicity scores derived from the persistence diagrams; higher (closer to 1) is better.

This work makes four key contributions. First, we introduce a novel, interpretable pipeline for automatically quantifying SMM in autistic individuals using sliding window persistent homology applied to time-series data derived from accelerometer sensors and video-based pose trajectories extracted from MediaPipe [24] (see Figures 1 and 2). Second, when applied to accelerometer data, we achieve classification performance on par with state-of-the-art frequency domain deep learning models that utilize simpler, more interpretable classifiers and features (see Table 2). Third, we demonstrate that even when applied to noisier, low-resolution video data, our method remains competitive, highlighting its robustness (see Tables 1, 2 and 3). Finally, we show that the resulting topological scores provide a quantifiable and interpretable representation of recurrent movement patterns that generalizes across individuals (i.e., person-agnostic) and modalities (see Figures 3 and 4). These collective findings suggest that persistent homology offers an interpretable and robust framework for SMM quantification in autism, with utility across high-quality sensor data and low-quality video recordings.

**Table 1:** Classification performance of Random Forest models using *PS*_10_ periodicity scores (Equation 13) derived from pose estimation and accelerometer data. Metrics are reported for binary (None vs. SMM) and multiclass (None, Rock, Flap, Flap-Rock) classification settings. Values reflect per-class precision, recall, and F1-score, along with the overall weighted F1-score.

| Class | Pose Estimation |  |  |  | Accelerometer |  |  |  |
| --- | --- | --- | --- | --- | --- | --- | --- | --- |
|  | Precision | Recall | F1 | Support | Precision | Recall | F1 | Support |
| *Binary Classification* |  |  |  |  |  |  |  |  |
| None | 0.83 | 0.95 | 0.88 | 449 | 0.89 | 0.95 | 0.92 | 511 |
| SMM | 0.91 | 0.72 | 0.80 | 319 | 0.93 | 0.86 | 0.89 | 396 |
| **Weighted F1** |  |  |  | **0.85** |  |  |  | **0.91** |
| *Multiclass Classification* |  |  |  |  |  |  |  |  |
| None | 0.80 | 0.92 | 0.86 | 449 | 0.89 | 0.94 | 0.91 | 511 |
| Rock | 0.87 | 0.78 | 0.82 | 259 | 0.93 | 0.86 | 0.89 | 299 |
| Flap | 0.22 | 0.09 | 0.13 | 43 | 0.68 | 0.72 | 0.70 | 65 |
| Flap-Rock | 0.50 | 0.12 | 0.19 | 17 | 0.75 | 0.56 | 0.64 | 32 |
| **Weighted F1** |  |  |  | **0.79** |  |  |  | **0.88** |

**Table 2:**
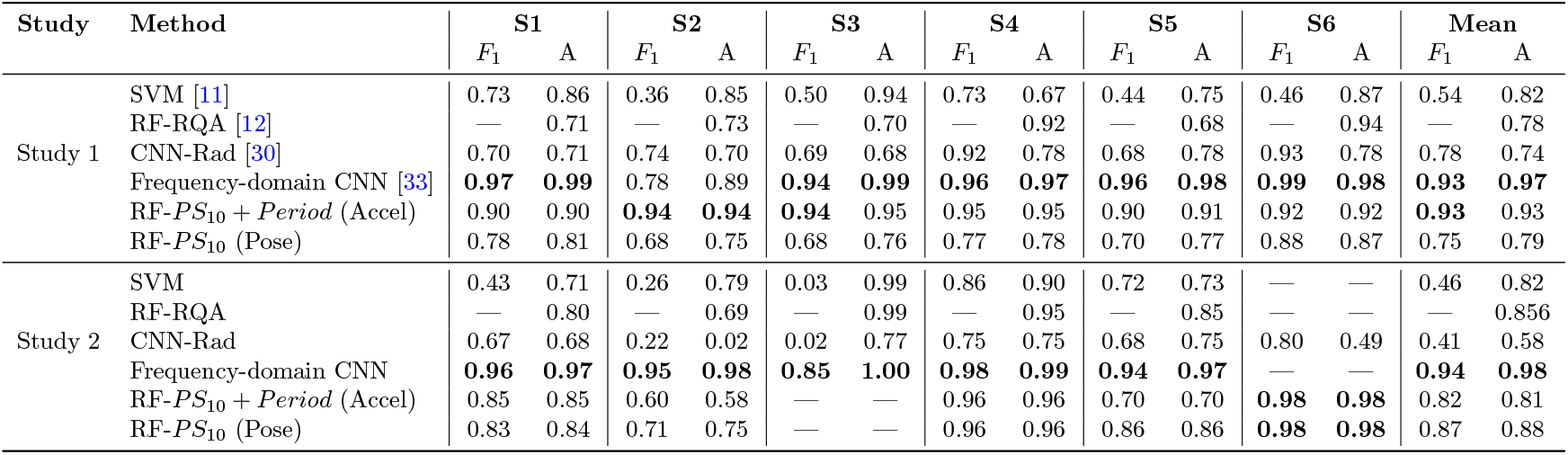
Comparison of our AQSM-*S*𝕎1*PerS* model trained on accelerometer data with additional period estimation features (RF-*PS*_10_ + *Period* (Accel)) and pose skeleton (RF-*PS*_10_ (Pose)) using a Random Forest classifier against prior work. Performance is evaluated using Accuracy (A) and weighted *F*_1_ score, the latter being more informative under class imbalance.

| Study | Method | S1 |  | S2 |  | S3 |  | S4 |  | S5 |  | S6 |  | Mean |  |
| --- | --- | --- | --- | --- | --- | --- | --- | --- | --- | --- | --- | --- | --- | --- | --- |
| | | $F_1$ | A | $F_1$ | A | $F_1$ | A | $F_1$ | A | $F_1$ | A | $F_1$ | A | $F_1$ | A |
| Study 1 | SVM [11] | 0.73 | 0.86 | 0.36 | 0.85 | 0.50 | 0.94 | 0.73 | 0.67 | 0.44 | 0.75 | 0.46 | 0.87 | 0.54 | 0.82 |
|  | RF-RQA [12] | — | 0.71 | — | 0.73 | — | 0.70 | — | 0.92 | — | 0.68 | — | 0.94 | — | 0.78 |
|  | CNN-Rad [30] | 0.70 | 0.71 | 0.74 | 0.70 | 0.69 | 0.68 | 0.92 | 0.78 | 0.68 | 0.78 | 0.93 | 0.78 | 0.78 | 0.74 |
|  | Frequency-domain CNN [33] | **0.97** | **0.99** | 0.78 | 0.89 | **0.94** | **0.99** | **0.96** | **0.97** | **0.96** | **0.98** | **0.99** | **0.98** | **0.93** | **0.97** |
| | RF-$PS_{10} + Period$ (Accel) | 0.90 | 0.90 | **0.94** | **0.94** | **0.94** | 0.95 | 0.95 | 0.95 | 0.90 | 0.91 | 0.92 | 0.92 | **0.93** | 0.93 |
| | RF-$PS_{10}$ (Pose) | 0.78 | 0.81 | 0.68 | 0.75 | 0.68 | 0.76 | 0.77 | 0.78 | 0.70 | 0.77 | 0.88 | 0.87 | 0.75 | 0.79 |
| Study 2 | SVM | 0.43 | 0.71 | 0.26 | 0.79 | 0.03 | 0.99 | 0.86 | 0.90 | 0.72 | 0.73 | — | — | 0.46 | 0.82 |
|  | RF-RQA | — | 0.80 | — | 0.69 | — | 0.99 | — | 0.95 | — | 0.85 | — | — | — | 0.856 |
|  | CNN-Rad | 0.67 | 0.68 | 0.22 | 0.02 | 0.02 | 0.77 | 0.75 | 0.75 | 0.68 | 0.75 | 0.80 | 0.49 | 0.41 | 0.58 |
|  | Frequency-domain CNN | **0.96** | **0.97** | **0.95** | **0.98** | **0.85** | **1.00** | **0.98** | **0.99** | **0.94** | **0.97** | — | — | **0.94** | **0.98** |
| | RF-$PS_{10} + Period$ (Accel) | 0.85 | 0.85 | 0.60 | 0.58 | — | — | 0.96 | 0.96 | 0.70 | 0.70 | **0.98** | **0.98** | 0.82 | 0.81 |
| | RF-$PS_{10}$ (Pose) | 0.83 | 0.84 | 0.71 | 0.75 | — | — | 0.96 | 0.96 | 0.86 | 0.86 | **0.98** | **0.98** | 0.87 | 0.88 |

**Table 3:** Binary classification performance (Weighted *F*_1_ Scores) of pose trajectories vs. accelerometer signals per participant under Leave-One-Individual-Out evaluation.

| Participant | Pose |  | Accelerometer |  |
| --- | --- | --- | --- | --- |
| | $PS_1$ | $PS_{10}$ | $PS_1$ | $PS_{10}$ |
| 1 | 0.81 | 0.81 | 0.83 | 0.85 |
| 2 | 0.71 | 0.71 | 0.68 | 0.68 |
| 3 | 0.72 | 0.68 | 0.92 | 0.94 |
| 4 | 0.89 | 0.88 | 0.92 | 0.96 |
| 5 | 0.79 | 0.79 | 0.74 | 0.82 |
| 6 | 0.89 | 0.91 | 0.94 | 0.93 |
| **Mean** | **0.80** | **0.80** | **0.84** | **0.86** |

**Figure 2:**
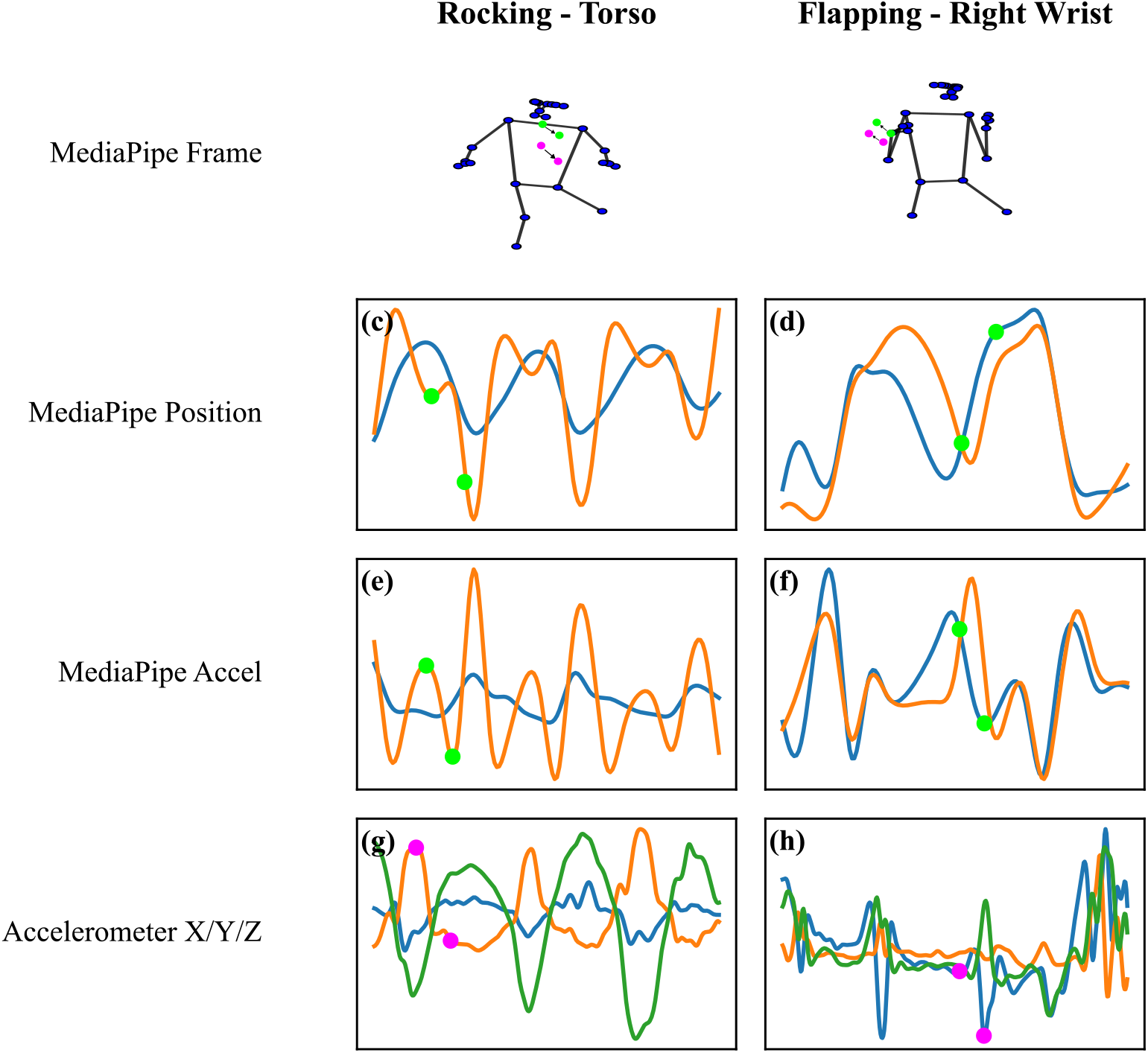
Multimodal comparison of autistic SMM across pose landmarks and accelerometer sensors. Columns represent distinct behaviors: Left, rocking involving torso motion; Right, flapping involving right wrist motion. Each row corresponds to a different data modality. (a, b): Video frames with MediaPipe skeleton overlays. Arrows indicate the direction of movement at a representative moment. Green dots denote the position and movement of a MediaPipe landmark, corresponding to matched timepoints in the time series plots. Purple dots indicate the position and movement of the accelerometer device at the same matched timepoints. (c, d): 2D trajectories of the chest (c), estimated from the midpoint of the shoulders and head, and the right wrist (d) landmarks from MediaPipe. The two green dots on each plot correspond to the same time points in the video frames above. (e, f): Approximated accelerations computed via second-order central differences, as defined in Equation 1. These estimates are sensitive to tracking noise and may not reflect absolute physical acceleration. (g, h): Tri-axial accelerometer signals from wearable sensors placed on the torso (g) and right wrist (h). All signals are consistently colored by axis: X (blue), Y (orange), Z (green). Purple dots correspond to the same timepoints in the video frames, illustrating temporal alignment across sensing modalities. Despite differences in sensing methods, coordinate frames, and signal units, periodic SMM is consistently captured. This visual correspondence supports using video-based pose estimation as a proxy for wearable inertial sensing in quantifying repetitive motor behavior.

**Figure 3:**
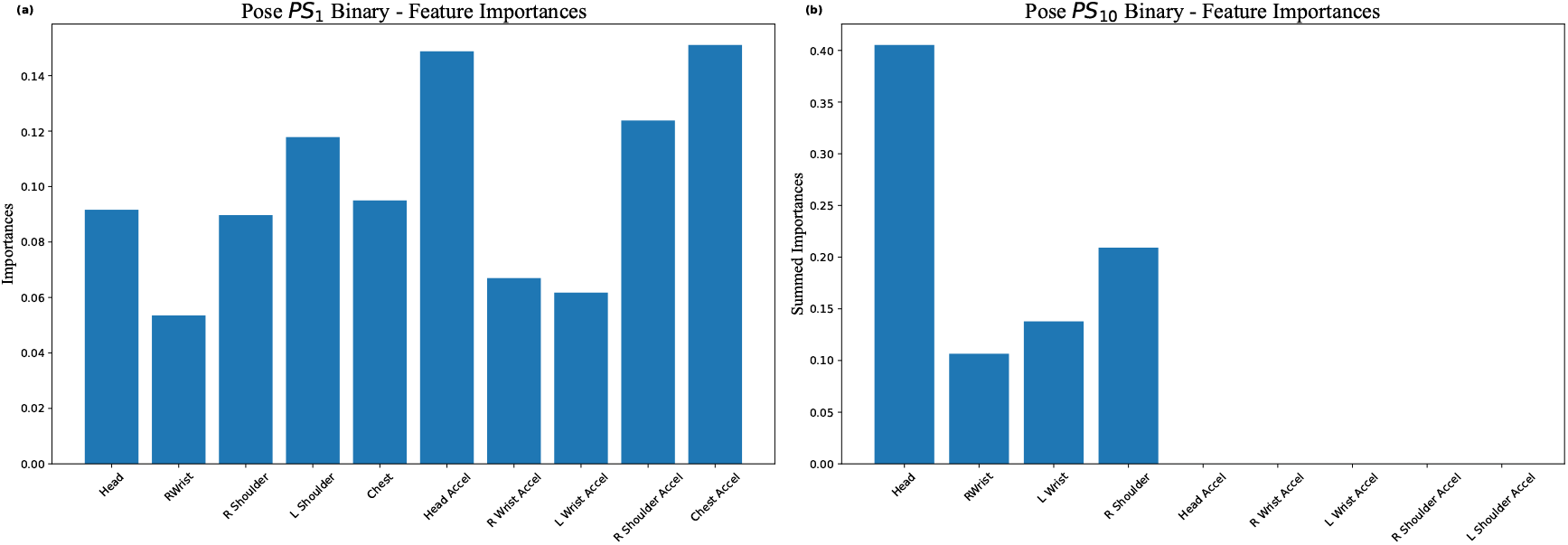
Feature importance for binary classification using pose-derived landmark position and derived acceleration features. In both the *PS*_1_ and *PS*_10_ models, features related to the torso and head contribute most significantly to the accurate detection of autistic SMM. (a) For *PS*_1_, features from all torso-related trajectories dominate the model’s decisions. (b) For *PS*_10_, spatial trajectory features of the head contribute the most. Note that certain landmarks are omitted because Bayesian optimization identified them as non-informative and excluded them from the final feature set. The consistent prominence of torso-related features aligns with prior findings [12] and reinforces the predictive power of upper-body motion dynamics in identifying autistic SMM.

**Figure 4:**
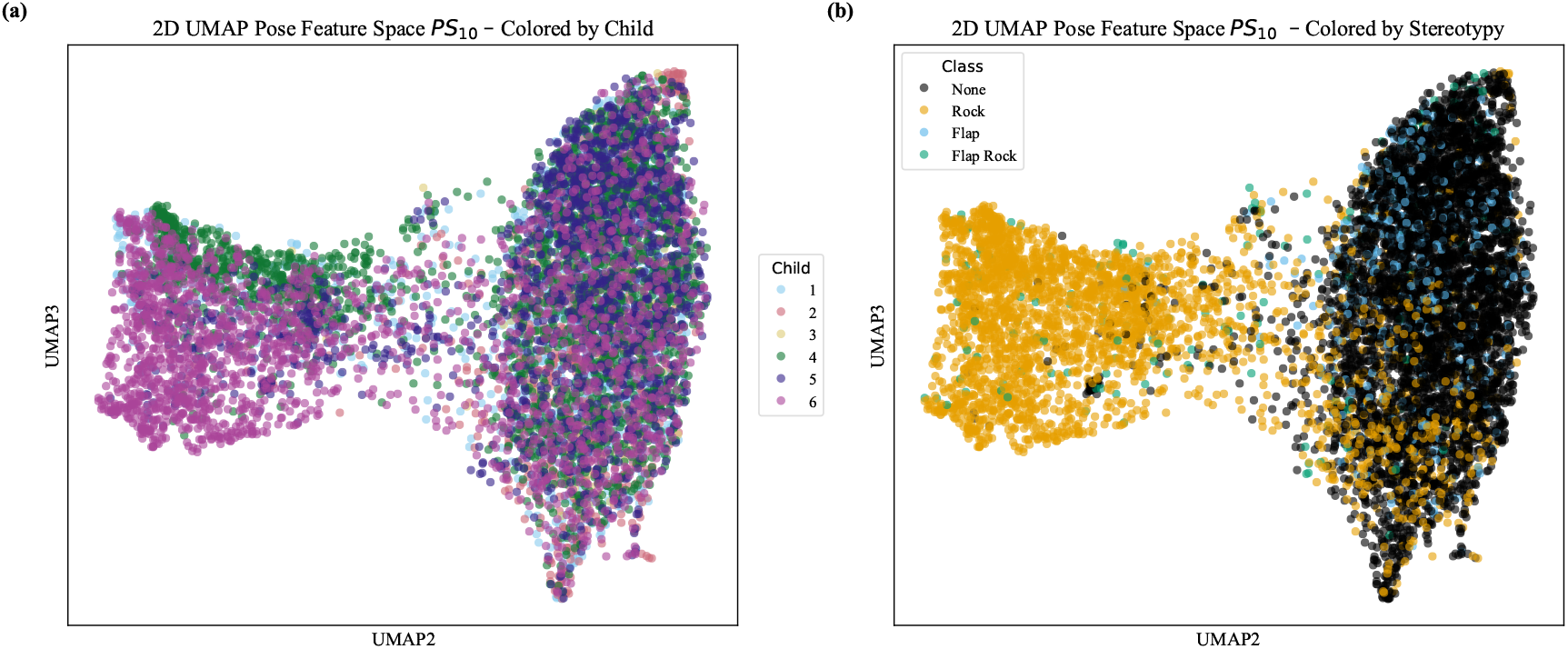
*UMAP* embeddings of pose-based landmark *PS*_10_ feature space colored by (a) participant ID and (b) SMM class. Note that the feature spaces are subsampled down for ease of visualization. In (a), the overlap across participants suggests no variability based on ID. In contrast, (b) shows clear organization based on SMM class, indicating that despite individual variability, our topological features encode behaviorally meaningful patterns that generalize across participants.

## Results

### Participants

We performed secondary analyses on a convenience sample of 6 children between 12 and 20 years from the Groden Center, RI (a school for children and adults with autism and other developmental difficulties) who participated in two published studies three years apart [1, 11]. The Groden Center human subjects review board approved the studies, and parental consent was obtained before participation. This included permission to release de-identified data to other data analysts for research purposes. Participants were included in the study if they met the following inclusion criteria: (a) had a documented DSM-IV-TR diagnosis of Autism Spectrum Disorder (ASD) made by a licensed psychologist familiar with the individual [2]; (b) were between the ages of 12–20 years; (c) had a clinically significant score (i.e., “behavior occurs and is a severe problem”) on the Whole Body and/or Hand/Finger items included in the Stereotyped Behavior Subscale of the Repetitive Behavior Scale [20]; and (d) exhibited, on average, at least 10 hand flapping or body rocking incidents per hour.

### Equipment

As described more fully in [1], participants were video recorded, and the recordings were annotated offline. Time segments noting hand flapping, body rocking, and a combination of both (i.e., flaprock) were labeled by their start and end times. These annotations (with 90% agreement between two independent raters) served as ground truth labels for our classification. The video recordings are approximately 30 minutes long and 4-6 fps. In addition to video data, participants wore three time-synchronized 3-axis accelerometers on each wrist and torso in both studies. Different accelerometers were used in each study. The first study employed MITes 3-axis sensors recording ±2 g data at 60 Hz. The second study utilized Wockets 3-axis sensors recording ±4 g data at 90 Hz. Recordings were made across multiple sessions within each study. These annotated videos and acceleration records served as signals with varying temporal resolutions to test and benchmark our data analysis pipeline.

### Setting, Procedure, & Data Obtained

All data collection sessions were conducted first in a laboratory setting and subsequently in naturalistic classroom settings, which included a diverse set of stimuli, demands for shared attention, and the presence of other students. In the classroom setting, each participant was observed seated at their desk during regular school hours (9:00 a.m.–3:00 p.m.). Observations included typical classroom activities (eating lunch, spelling program, sorting, etc.), with participants working independently and with a familiar teacher present. Observed SMM across both settings was naturally emitted, not experimentally elicited or consequently intervened upon.

### Temporal Segmentation

We segmented recordings into overlapping time blocks to analyze the periodicity of movements as a proxy for the presence of SMM. To find the optimal block size, we computed the following mean durations for each type of SMM:

1. Rock - 11.78 seconds
2. Flap - 5.26 seconds
3. Flap-Rock - 4.00 seconds

Supplementary Table 1 and Supplementary Figure 1 present descriptive statistics on the number and duration of flap, rock, and flap-rock bouts recorded and a visual depiction of their duration distributions. A 4-second window size was chosen to ensure sufficient time to capture at least two motion oscillations, but not so large as to miss SMMs with shorter durations. The step size is variable depending on the experiment performed.

### AQSM - *S*𝕎1*PerS* Feature Extraction

Using time series data as input, we analyzed signals from accelerometers and pose trajectories extracted from pre-recorded videos. For the video modality, MediaPipe provided spatial trajectories of key landmarks (e.g., head, wrists, shoulders), and we estimated a “Chest” point as the midpoint between the shoulders and head to obtain a more stable reference. Using these pose trajectories, acceleration was computed as an additional representation of movement. We applied the following pipeline to 4-second time intervals across all recordings (see Methods Section for more details):

1. *Pre-process Time Series*: We smoothed the raw spatial trajectories of each landmark, resulting in 2-dimensional time series *F*_*landmark*_(*t*) = (*f*_*x*_(*t*), *f*_*y*_(*t*)). Additionally, we used the acceleration estimate of *F*_*landmark*_ as an alternative representation of movement, providing further smoothing. This results in 6 landmarks with two motion representations, totaling 12 individual time series. For accelerometer data, we used the 3-dimensional sensor readings located on the torso, left wrist, and right wrist *F*_*sensor*_ = (*f*_*x*_(*t*), *f*_*y*_(*t*), *f*_*z*_(*t*)), analyzing a total of 3 time series.
2. *Data Interpolation*: To compensate for the videos’ low frame rate (i.e., there are about 20 time points in a 4-second window), we interpolated 1000 values using cubic splines. This higher resolution allows for more accurate period estimation. To ensure an equal sampling rate for the accelerometer data, each sensor was resampled to 95Hz.
3. *Remove Linear Trends*: We removed linear trends caused by gradual camera drift and global motion from the time series because they can distort and obscure periodic patterns and sliding window embeddings.
4. *Estimate Period*: We determined the period *L* via the Complex Discrete Fourier Transform to obtain the optimal parameters *τ >* 0 and *d* ∈ ℕ for the sliding window embedding *SW*_*d,τ*_ *F* of the pre-processed landmark time series *F*. Methods for estimating the period of accelerometer data can be found in Supplementary Note 4.
5. *Populate Sliding Window Point Cloud*: We evaluate *X* = {*SW*_*d,τ*_ *F* (*t*): *t* ∈ *I*} ⊂ ℝ^48^ where *I* is a finite set of time values.
6. *Normalize Point Cloud*: We apply the *k*-nearest neighbor density filter to only capture meaningful data. Then, we apply centering and sphere normalization to enable comparisons of movements between different people, controlling for different amplitudes, and providing a way to manage signal dampening.
7. *Compute Score*: We compute the persistence diagram of the normalized sliding window point cloud and calculate the periodicity score (either *PS*_1_ or *PS*_10_) from features with the largest persistence. Our implementation uses the library Ripser.py [35].

### Experimental Results

The following experimental results are based on the feature extraction and model optimization schemes described in the *Methods* section. For reference, the model used for classification is a Random Forest, known for its scalability, high-dimensional data handling, and ability to discern which features contribute the most to classification. We partitioned the set of feature vectors in each experiment into training, testing, and validation sets to optimize and evaluate model performance and quantify generalizability across unseen data.

#### Experiment 1 - Stratified Set Classification

In our first experiment, we assess the effectiveness of our video-based pipeline in binary (SMM vs None) and multi-class (Rock vs Flap vs Flap-Rock vs None) classification compared to wearable accelerometer data. To this end, the data is split into training, testing, and validation subsets, ensuring equal behavioral variability. Specifically, we sought a proportional distribution of classes from each participant’s recordings to ensure that the classifier does not become biased towards movements exhibited by a specific individual. We used *scikit-learn*’s `train_test_split` to split data into subsets randomly while respecting class distributions. We then aggregated these splits to form the final training, test, and validation sets. This approach ensures fair representation of all participants and class labels across subsets. This experiment has utility in two areas: (1) evaluating feature importance in scenarios where both individual variability and class imbalance must be accounted for; and (2) ensuring that all participants are represented in all splits, enabling the evaluation of model behavior across a wider diversity of data, and allowing better insight into when and how the model fails. Another consideration is data leakage, i.e., the potential for overlapping time intervals of the same participants and sessions in multiple subsets. In this case, the model may encounter nearly identical information during training and testing, which can artificially inflate its evaluated performance. To overcome this challenge, we chose a step size of 4 seconds to eliminate overlap between segmented video intervals.

Table 1 summarizes results from Experiment 1 using pose trajectories and accelerometer data for binary and multi-class classification. For ease, this table only reports *PS*_10_ scores as they yield better accuracy; refer to Supplementary Table 2 for a full table.

We investigated the lower performance in multi-class SMM classification, specifically flapping and flap-rock, by manually reviewing all 4-second windows labeled accordingly. Pose estimation included visual inspection of the MediaPipe pose skeleton movement and the derived wrist time series to assess the true periodic nature of the movement and whether implementation issues could explain observed results. We performed this analysis twice and found that only approximately 12% of flapping instances clearly exhibited periodic motion. Reasons for this include: (1) occlusions causing the skeleton not to track the motion correctly, or (2) subtle finger movements or hand tremors annotated as flapping, which MediaPipe pose cannot track. These observations highlight a fundamental challenge in modeling stereotypical flapping behaviors in autistic individuals. While higher-resolution video data and annotations would strengthen our analysis, collecting them poses challenges. There is limited availability of publicly available, naturalistic autism video datasets that include clinical and demographic metadata. Moreover, obtaining precise annotations for stereotypical behaviors like flapping or rocking is time-intensive and often subjective, requiring expert raters and substantial inter-rater agreement. These constraints make it difficult to produce “ideal” data at scale. Therefore, we opted to work with the best available real-world data from previously published studies where autism experts annotated videos.

SMM are typically repetitive, rhythmic patterns sustained over extended durations that result in consistently periodic behavior. Hand flapping tends to exhibit greater frequency, amplitude, and duration variability, and occurs in brief, rapid bursts, leading to less consistent periodic behavior [1]. Our model’s performance with pose- and accelerometer-based methods supports this distinction (See Supplementary Figs. 4–15 and Supplementary Table 2), where rocking-related movements consistently produced high accuracy (Average Precision ≥ 87%). Conversely, flapping-related motions yielded lower performance (Average Precision ≤ 81%) and were confused with non-stereotypical behavior. Using accelerometer data with high temporal rates yielded higher results than our pose landmarks, as it captures higher-frequency oscillations. While “flapping” includes rhythmic, repetitive motion of the hands, these behaviors often manifest with variable intensity and frequency, wherein accurate “ground-truth” differs between annotators. Periodicity is a useful and frequently present feature of many SMMs, particularly hand-flapping and rocking. Still, it is not a defining characteristic in clinical or research settings. Instead, repetition and stereotypy are broader terms that allow for rhythm, frequency, and duration variation. Accordingly, we use periodicity as an informative but non-exclusive feature in our analysis that accounts for most annotations in our dataset. Since the accelerometer data has a high sampling rate, we included the estimated period as one frequency-domain feature along with our periodicity score. Period estimation, already computed within the *S*𝕎1*PerS* algorithm, is included as a feature because it directly captures the repetition rate of SMM. Given the low sampling rate of the video data, the period feature was not informative in the model and thus not included. Supplementary Table 2 shows the increase in classification accuracy when using this simple, interpretable feature.

Figure 3 illustrates which topologically-derived features contributed most to the Random Forest’s decisions using pose-based landmarks derived from video data. The head, wrists (L Wrist and R Wrist), shoulders (L Shoulder and R Shoulder), and chest landmark trajectories, along with their second temporal derivatives (Accel), were analyzed as described in the *Methods* section. The bars indicate the contribution of each feature to the model. The feature importance measure of the Random Forest classifier indicates that the wrist signals contributed less discriminative power than the torso-related landmarks, as expected, due to the challenges articulated with flapping-related movements. Refer to Supplementary Figs. 16-17 for feature importances of the binary model using accelerometer sensors instead of pose-based landmarks.

#### Experiment 2 - Leave-One-Session-Out

To enable closer comparison with established baselines in the field that evaluate binary classification performance per session [11, 12, 30, 33], we adapted our pipeline to train and evaluate models on a per-session basis. The prior models were all trained on the 3-axis accelerometer time series data provided in the same datasets used in the present study. Our pipeline’s *S*W1*PerS* component is the key topological method for computing periodicity scores (see *Methods*). We applied them to the accelerometer signals in the same dataset used in prior studies (see Supplementary Note 4). This enabled us to extract topologically-derived scores from the same input modality, allowing for a more consistent comparison across approaches. We split the data so that videos from the same participant in a study are used for testing, and the remaining data is split 80/20 for training and validation.

Table 2 summarizes results from Experiment 2 using *PS*_10_ and the pose and accelerometer data for binary classification. We evaluate the performance of our methods against other SMM detection methods, such as Support Vector Machines (SVMs) [11], Random Forest with Recurrence Quantification Analysis (RF-RQA) [12], and deep learning approaches based on Convolutional Neural Networks (CNNs) [30, 33]. A key methodological difference between our method and prior work is that our pipeline requires each window to contain at least four seconds of annotated SMM to capture meaningful structure, as this allows at least two oscillations of movement. As a result, shorter duration SMM is not analyzed. Prior methods use 1-second segments with 87% overlap compared to our 4-second segments with ~ 95% overlap (step size of 1*/fps*). While these differences preclude strict one-to-one comparability, they offer valuable insight into the tradeoffs between model complexity and generalization across different analyses. In Table 2, we present our best model that computes *PS*_10_ features along with the period estimates on the accelerometer data. For details of *PS*_1_ comparison, please refer to Supplementary Table 3. While the frequency-domain CNN [33] achieved the highest performance across most sessions, our model offers comparable performance with the added simplicity and interpretability of a Random Forest classifier and our mathematically principled features.

#### Experiment 3 - Leave-One-Individual-Out

This experiment tests how well TDA features generalize to truly unseen data. We adapt Leave-One-Session-Out, where the model is tested on one person’s data, and data from the remaining people is used for training and validation, along with Bayesian optimization to find near-optimal hyperparameters. Feature extraction remains the same as Experiment 2. Leaving an individual out during evaluation, regardless of session, offers a more rigorous assessment of generalization and better reflects real-world deployment scenarios where models are applied to completely unseen people.

Table 3 summarizes the results from Experiment 3 using both *PS*_1_ and *PS*_10_ with pose and accelerometer data for binary classification. Supplementary Table 4 includes the performance of the period features with accelerometer data. As expected, classification performance with pose data was lower for participants 2 and 3, who exhibited higher frequency flapping behaviors involving occlusions or fine hand movements. MediaPipe finds such movements near impossible to track accurately. In contrast, using accelerometer data achieved higher accuracy for these participants, likely due to its sensitivity to subtle motion patterns irrespective of visibility.

#### Experiment 4 - Feature Space

Using our pipeline at 4-second intervals with a step size of 1*/fps*, as in Experiments 2 and 3, we employed Uniform Manifold Approximation and Projection (*UMAP*) to visualize our feature space. *UMAP* is a method for dimensionality-reduction that finds a low-dimensional representation of high-dimensional data while preserving topological structure [26]. We used it to observe how our feature space using pose landmarks, which is 12-dimensional using *PS*_1_ or 120-dimensional using *PS*_10_, is organized by coloring each point, representing the data of a single 4-second time interval, based on participant or SMM.

Figure 4 illustrates that our model is person-agnostic, meaning that regardless of which participant the data came from, points corresponding to the same behavioral class tend to occupy overlapping regions in the feature space. This suggests our model captures generalizable features of SMM and is blind to person-specific mannerisms. The feature space organization from both pose-estimation and accelerometer readings using *PS*_1_ and *PS*_10_ scores (see Supplementary Figs. 18-20) also demonstrates that the feature space is person-agnostic and that flapping-related motions are mixed with no SMM when using pose landmarks. When looking at the accelerometer feature space visualizations (Supplementary Figs. 19-20), it is clearer that there is some distinction between flapping and no SMM.

### Application of Methods

To complement the classification and visualization analyses presented above, we applied our individual participant-level methods to demonstrate its interpretability and potential clinical utility.

#### Within-Person Illustrations

Supplementary Figures 23 and 24 provide an example of median periodicity score trajectories for a single participant across multiple sessions, stratified by annotation (None, Rock, Flap, Flap-Rock). Supplementary Figure 21 summarizes *PS*_1_ scores computed on pose landmark trajectories (chest, left wrist, right wrist, and their acceleration estimates), aggregated as session-wise medians within each annotation. Panels (c) and (f) indicate that the topological strength of repetition increases for rocking-related movements for this participant. The same trend is shown in Supplementary Figure 24, which reports the corresponding median persistence trajectories computed from accelerometer sensors (torso, left wrist, right wrist), again stratified by annotation and session. Panel (a) of Supplementary Figure 22 demonstrates that the strength of repetition for the torso sensor increases for rocking-related movements. These observations suggest that the individual’s stereotypy expression is intensifying (i.e., stronger, more periodic patterns).

These visualizations provide a concrete demonstration of how median *PS*_1_ scores fluctuate both over time and across behaviors within the same individual.

#### Summary of Periodicity Scores

To highlight the practical utility of our features, Supplementary Tables 4-7 report the median ± SD of *PS*_1_ scores across participants, sessions, and annotations for a representative set of landmarks (chest, left wrist, right wrist, and their acceleration estimates) and the accelerometer sensors (torso, left wrist, right wrist). Supplementary Tables 5 and 6 report the median ± SD *PS*_1_ scores per pose landmark stratified by participant, session, and annotation. In the laboratory setting (Study 1), according to the median *PS*_1_ scores, the participant’s rocking behavior is localized primarily to the torso. In Study 2, conducted a few years later in a naturalistic classroom environment, rocking constitutes a broader full-body movement involving both the torso and arms. The repetition strength (*PS*_1_ score) also increases from Study 1 (lab) to Study 2 (classroom). Supplementary Tables 7 and 8 present the same report using accelerometer data, reinforcing the observation that the participant transitions from localized rocking in Study 1 to more full-body rocking in Study 2, with corresponding increases in repetition strength. Full-body rocking and increased repetition strength from Study 1 to Study 2 may reflect increased intensity of self-regulatory behavior or developmental changes in the expression of SMM. Context differences are also relevant. The controlled laboratory setting may constrain the expression of rocking. In contrast, the more socially active classroom setting may prompt whole-body movements that are more repetitive. We observe this trend not only in participant 1 but also in participants 2, 4, and 5, though at varying degrees. This suggests a potential link between time, setting, topography, and intensity of autistic SMM.

#### Summary of Period Estimates

Finally, a key part of our methodology is estimating the period of the motion time series, which we report alongside periodicity scores to provide an additional temporal marker of stereotyped movement. While persistent homology quantifies the strength of repetition, period estimates capture the dominant rhythmicity of the movement and thereby complement persistence in distinguishing between annotations and across participants. Supplementary Tables 8-11 summarize period estimates as median [Inter-Quartile-Range] stratified by participant and annotation. To estimate the period of movements, we restricted the analysis to a physiologically-plausible frequency range for SMM contained within a 4-second time window (0.25-6.0 Hz - periods between 0.17 and 4 seconds). We note that the low frame rate of the video data limits our ability to estimate the period of high-frequency movements. Consequently, the analysis of median [Inter-Quartile-Range] estimated periods from the pose landmarks is less informative than the accelerometer data. This further underscores the need for high-quality video data. As a proof-of-concept, we demonstrate with the accelerometer data (Supplementary Tables 11 and 12) what could be achieved if similarly fine-grained temporal information were available from video.

As in the example above, we observe a similar trend in the median period estimations from the laboratory study compared to the classroom study. Across participants, we observe that median estimated periods consistently decrease from Study 1 to Study 2 across all sensors (Torso, Left Wrist, and Right Wrist). This indicates an increase in the frequency of repetitive movements over time. For example, Participant 1’s torso median period decreases across sessions during rocking and flap-rock - an inverse relationship to the periodicity scores. Consistent with the behavioral observation of more vigorous, full-body rocking in Study 2, the accelerometer data reveal increased frequency of torso movement along with wrist movement during rocking bouts, as evidenced by lower estimated median periods in the torso and wrists. A similar trend is again observed for Participants 2, 4, and 5, whose elevated *PS*_1_ scores in Study 2 during rocking are accompanied by shorter torso median periods. These observations suggest that SMM is faster and more repetitive in the classroom. Similar to the median *PS*_1_ scores, changes in motor stereotypy expression could be caused by environmental differences that impose different regulatory demands or reflect developmental progression over the three-year interval.

## Discussion

We introduce a novel pipeline that integrates time series–based motion analysis (from both computer vision–derived pose trajectories and wearable accelerometers) with persistent homology to recognize and quantify recurrent motor behaviors. Our primary application is the automated quantification of autistic SMM. We developed an interpretable set of topological features which, when applied to accelerometer data with simple classifiers, achieved performance comparable to that of complex neural networks, and remain competitive when applied to pose landmarks extracted from noisy, low-quality video recordings. The person-agnostic design of our method across pose and accelerometer data supports broader generalization and may enable large-scale, clinically informative analyses of SMM. For example, time series analysis of our model’s parameters could reveal patterns in movement intensity or variability within individuals over time. Survival analysis and point process models of our parameters could estimate the probability of SMM, compare rates across different movements and individuals, and examine how factors like environment or physiology influence these rates. Pattern-based approaches, such as cluster analysis and growth mixture models, using our parameters could categorize individuals into groups with similar or different movement types and frequencies, potentially uncovering underlying biological mechanisms and environmental factors that sustain these behaviors [25]. Consistent movement rates across environments could imply a biological basis, while fluctuations in response to stimuli or settings could indicate behavioral, sensory, or homeostatic influences [32]. Such analyses could also inform and assess interventions when deemed necessary [31], offering predictions about the most effective strategies [32]. For example, individuals with high stereotypical movements across all environments might benefit from medication [22], while those whose behaviors are environment-specific might respond better to behavioral therapies [6, 21, 14]. Finally, MediaPipe is cross-platform and lightweight, facilitating wider-scale comparisons of SMM in autism and other neurodevelopmental conditions (Rett’s Syndrome, Angelman’s syndrome, Tourette’s, etc.) where repetitive motor movements are a primary feature.

While our analysis suggests that periodicity is a meaningful and biologically interpretable characteristic of SMM, it is not a sufficient descriptor to capture their full variability. Accordingly, we frame periodicity as a starting point for analysis rather than a comprehensive representation of repetitive motor behavior. Future work could explore other features that more robustly capture the spectrum of SMM. For example, simply including the period estimates boosted classification power with the accelerometer signals. Future research could also strive to develop a reliable way to track changes in depth for a more holistic representation of SMM. Using multiple cameras in the same room may be a good way to accomplish this. Applying our method to a larger data set with improved frame rates from a wider sample of autistic individuals is also required to address differences in sex, age, and race. Finally, a more robust way of tracking fine hand and finger movements is needed. While MediaPipe supports a holistic landmark solution that outputs 33 pose landmarks, 468 face landmarks, and 21 landmarks per hand, the low quality of our (and most existing) video data resulted in unreliable and inaccurate detection of hand landmarks. Higher frame-rate videos would enable the capture of faster hand movements, particularly those occurring at higher frequencies. This is evident from the accelerometer data, which we used as a control to illustrate the upper bound of motion frequencies that can be detected and how well our pipeline can perform on high temporal-resolution data.

## Methods

### MediaPipe Pose Estimation + YOLO Child Tracking

MediaPipe [24] is an open-source suite developed by Google, implementing efficient, cross-platform solutions for real-time machine learning pipelines. We used MediaPipe’s pose land-marker tracking to analyze full-body poses and actions. Given a video frame showing one individual, MediaPipe outputs the spatial (*x, y*) positions of 33 landmarks using the BlazePose [5] model. Like other pose detection models, MediaPipe can struggle to detect the correct person, so we utilized YOLOv5 [16] as a means to track the participant under observation throughout a video. The YOLO model outputs a bounding box around the desired individual frame by frame. For each frame, MediaPipe is applied within the bounding box to track the pose of the correct person; see Figure 5 for an example. Training details of the YOLO model are provided in Supplementary Note 2, and our full implementation can be found in the GitHub repository replicating the results of this paper https://github.com/ambaye15/AQSM_SW1PerS.git.

**Figure 5:**
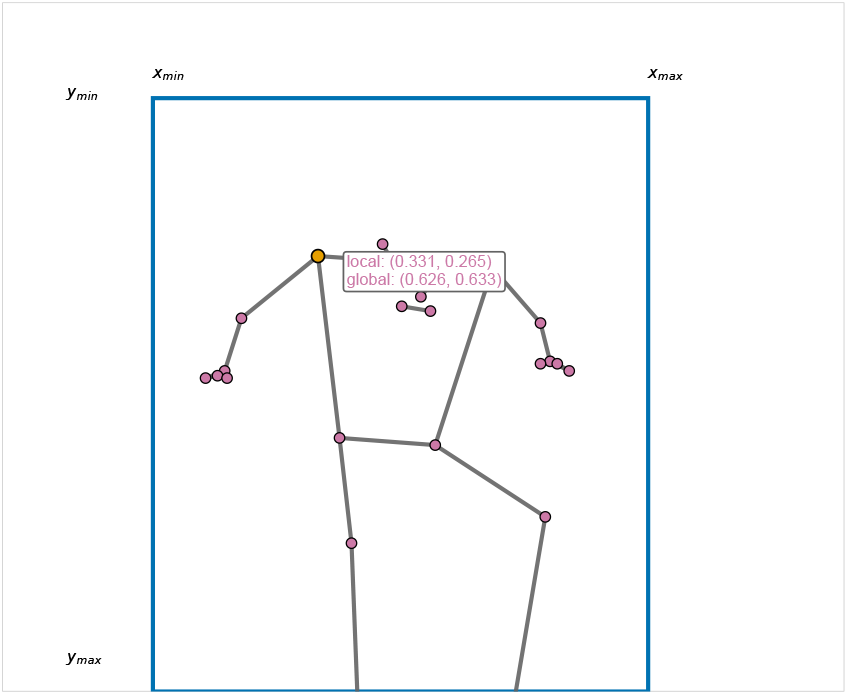
Visualization of pose estimation and bounding box tracking. The YOLOv5 bounding box (blue rectangle) detects the person of interest, while the MediaPipe skeleton (gray) and landmarks (magenta) are estimated within the cropped region. The right shoulder joint is highlighted (yellow dot), with both local (normalized within the cropped region) and global (normalized to the full frame) coordinates annotated. When moving, both the landmarks and the bounding box may move simultaneously, resulting in either a constant position or loss of absolute motion when observing the local coordinates. Hence, global coordinates are necessary to capture the full dynamics of movement. Note that in the transformation from local to global coordinates, the x and y axes are mirrored, which does not affect periodicity analysis as long as the transformation is applied consistently.

### YOLO Box Tracking and Smoothing

The YOLO model produces bounding boxes frame by frame, which can experience temporal jitter and sharp movements due to noise, occlusions, or abrupt motion. The output bounding boxes for the YOLO model are in the form (*x*_*min*_, *x*_*max*_, *y*_*min*_, *y*_*max*_). Because MediaPipe is used within each bounding box, sudden jumps or shrinkages can introduce substantial noise or result in incomplete pose landmarks, negatively affecting downstream analysis. To reduce the noise of bounding box predictions, we implemented a Kalman filter [3, Section 3.1] that predicts the position of the bounding box for a given frame based on the previous frames and the latest YOLO prediction. Indeed, let

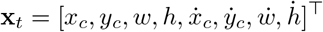

be the state column vector that models the position and velocity for the bounding box center, width, and height. The state transition model predicts the next state at time *t* + Δ*t*, for Δ*t >* 0, as:

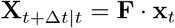

with

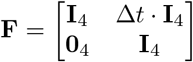

being the state transition matrix that encodes constant-velocity motion, where the new position is updated as:

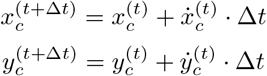

We use the `KalmanFilter` module from *FilterPy* to implement.

### Landmark Extraction

Given the estimated bounding box from YOLO, MediaPipe computes the position of each pose landmark within that box. Since these landmark coordinates *x, y* ∈ [0.0, 1.0] are local to the bounding box, whose coordinates in the frame are (*x*_*min*_, *x*_*max*_, *y*_*min*_, *y*_*max*_) (where, e.g. *x* = 0 in landmark coordinates is *x*_*min*_ in the frame), we transform them into frame coordinates that are globally consistent among different frames. Given an overall frame dimension of *w* × *h*, this leads to the following transformation:

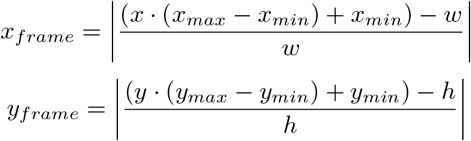

This transformation subtracts the unnormalized coordinate from the full frame width/height before normalizing, effectively shifting the origin from the top-left to the bottom-right. The absolute value then mirrors the coordinates back into the [0.0, 1.0] range, reversing the direction of both axes. That is, left becomes right and top becomes bottom. The result is a mirrored coordinate system. However, this does not affect periodicity analyses since the motion is consistently tracked in the whole frame and the transformation preserves relative displacements.

See Figure 5 for an example of a predicted YOLO bounding box, the MediaPipe skeleton fitted to the detected participant, and the local/global coordinates of one of the detected landmarks.

### Landmark Smoothing

Accurate temporal tracking of landmarks can be challenging for low-resolution video data, particularly for the wrists, which experience rapid, high-frequency flapping motions in the present context. MediaPipe can struggle to track these fast motions, causing the estimated wrist trajectory to exhibit inhuman movements. We detect these abrupt changes by calculating the acceleration of the landmarks. Specifically, given *f* (*t*_0_), …, *f* (*t*_*n*_) ∈ ℝ (e.g., the values of the *x* coordinate for a particular landmark) and Δ*t >* 0 we let:

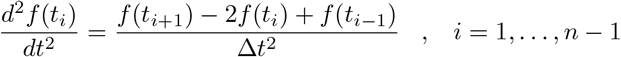

and for boundary points we use the second-order forward and backward derivatives, respectively:

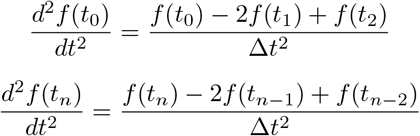

The resulting acceleration time series *a*(*t*) is thus:

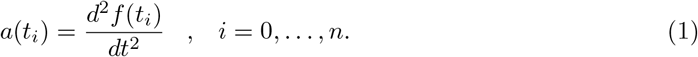

To flag unstable regions of wrist movement, we utilized the elbow flexion angles to detect when pose estimation fails. When wrist tracking is inaccurate, we observe that the elbow flexion angle, computed using the shoulder, elbow, and wrist coordinates

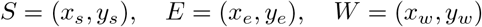

undergoes an abrupt change. Define the upper arm vector as

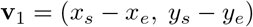

and the lower arm vector as

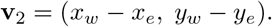

The elbow flexion angle *θ* at time *t* is given by

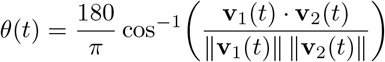

expressed in degrees, and we estimate the angular acceleration *a*(*θ*(*t*)) using second-order finite differences as before:

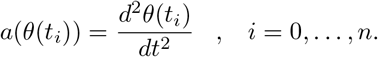

Analyzing angular acceleration patterns throughout each video, we chose a threshold of 500 degrees*/s*^2^ to maximize the number of flapping instances while filtering out noisy regions. Specifically, if

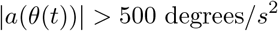

than the corresponding frame is deemed to have an unreliable pose estimate and is flagged for removal. In cases where the first or last frame is flagged as unreliable, we apply boundary extension, where the first valid value encountered is used to backfill the beginning, and the last valid value is used to forward fill the end. For internal frames, we use cubic interpolation to fill in missing gaps.

A similar filtering scheme is applied to the landmarks associated with rocking (i.e., the shoulders and head). By computing second-order finite differences for the *x* and *y* coordinates, we flag a frame as unreliable if either condition is met:

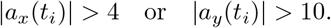

This approach minimizes the impact of pose estimation errors, such as jumping to a different person, by excluding outlier frames from further analysis [7].

Finally, we use a Gaussian filter to smooth the motion curves. The filter weights nearby points using a discrete Gaussian kernel derived from the continuous Gaussian function:

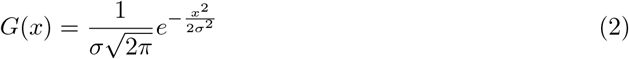

For a discrete signal *f*: {0, …, *T* − 1} → ℝ, the smoothed signal 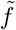 is given by:

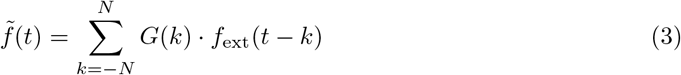

where *G*(*k*) is the discretized Gaussian kernel, *σ* controls the smoothing strength (we set *σ* = 0.5 for relatively low smoothing), and *f*_ext_ denotes the boundary-extended version of *f* (using e.g. reflection). For implementation, we use `gaussian_filter1d` from *scipy*.*ndimage*, where the default is *N* = 4*σ*. For a two-dimensional signal *F* (*t*) = (*f*_*x*_(*t*), *f*_*y*_(*t*)), we apply smoothing component-wise: 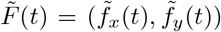. This reduces noise while preserving overall patterns in the motion curves.

### Sliding Window Embeddings

To recognize periodicity (e.g., in the movement of estimated landmark locations), one can transform the problem of recurrence detection in time series data into that of shape analysis, where tools from TDA can be applied. Specifically, we use the Sliding Window Embedding, which transforms a time series into a discretized point cloud in a high-dimensional space [28]. When the signal is periodic, the arrangement of the point cloud is circular. Formally:

#### Definition 1

(Sliding Window Embedding of *m*-Dimensional signals). *Let F*: ℝ −→ ℝ^*m*^ *be an m-dimensional time series, m* ≥ 1, *and write F* (*t*) = (*f*_1_(*t*), …, *f*_*m*_(*t*)). *Given parameters d* ∈ ℕ *and τ >* 0, *the sliding window embedding of the signal F at t* ∈ ℝ *is the matrix*

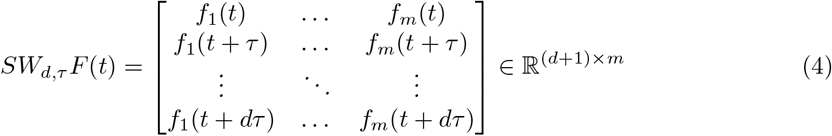

*Moreover, given a set of time values I* ⊂ ℝ, *the sliding window point cloud of F is the set X* = {*SW*_*d,τ*_ *F* (*t*): *t* ∈ *I*} ⊂ ℝ^(*d*+1)×*m*^. *We use the Euclidean (i*.*e*., *Frobenius) norm* ‖ · ‖ *in* ℝ^(*d*+1)×*m*^ *to measure pairwise distances* ‖*p* − *p*^′^‖ *between points p, p*^′^ ∈ *X*.

When the signal *F* is only observed at discrete time points *t*_0_ *< t*_1_ *<* · · · *< t*_*n*_ we extend it to *t* ∈ ℝ before computing its sliding window point cloud, as follows:

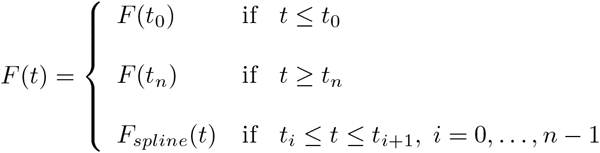

where *F*_*spline*_: ℝ −→ ℝ^*m*^ is the function whose *j*-th coordinate, *j* = 1, …, *m*, is the cubic spline interpolating the points (*t*_*i*_, *f*_*j*_(*t*_*i*_)) for *i* = 0, …, *n*. See Figure 7, panels (a) and (b), for an example with *m* = 2.

### Sliding Window Point Cloud Filtering and Normalization

Dense regions in the sliding window point cloud *X* ⊂ ℝ^(*d*+1)×*m*^ correspond to areas capturing the global shape and geometry of the object. More sparse regions, on the other hand, can contribute noise, degrading topological inference. We use a *k*-nearest neighbors density approach to filter out this sparse data. That is, for *k* ∈ ℕ and each *p* ∈ *X*, let *N*_*k*_(*p*) ⊂ *X* be the set comprised of the *k* nearest neighbors of *p* in *X* as measured by the Euclidean distance in ℝ^(*d*+1)×*m*^. To filter based on density, we estimate the density of a point as the reciprocal of the median distance to its *k*-nearest neighbors:

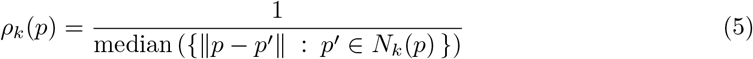

Given the values *ρ*_*k*_(*p*) for *p* ∈ *X*, we let the subsample of densest points be

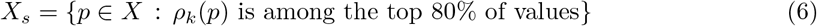

After subsampling the point cloud, we mean-center each point individually and normalize it to enable comparisons between periodic motions of different videos and varying ranges or intensities. That is:

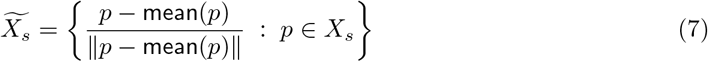

where `mean` (*p*) ∈ ℝ^(*d*+1)×*m*^ is the matrix with all entries equal to the average of the entries of the (*d* + 1) × *m* matrix *p* ∈ *X*_*s*_. Density thresholding in the sliding window point cloud ameliorates the impact of noisy recurrent behavior in *F*, and mean centering helps remove linear trends from the original signal. As a result, the point cloud 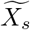 is a subset of the unit sphere in ℝ^(*d*+1)×*m*^, whose periodicity score computed from persistent homology with appropriate normalization (Equation 11), is a number between zero and one.

### Persistent Homology of Sliding Window Point Clouds

Persistent homology quantifies the underlying topology of a shape 𝕏 given a finite, discrete sample *X*. By topology, we mean properties invariant to continuous deformations, such as the number of connected components in 𝕏, or whether it has holes or voids [13]. This is relevant to our analysis because if a signal *F* is periodic, then its centered and normalized sliding window embedding (with appropriate *d* and *τ*) describes a circle, sampled by the centered and normalized sliding window point cloud.

We proceed as follows. Given a set of points *X* with pairwise distances *d*_*X*_: *X* ×*X* −→ [0, ∞), the *Rips complex* at scale *ϵ* ≥ 0 is the triangulated space (more formally, the simplicial complex) with one vertex for each *x* ∈ *X*, one edge {*x*_0_, *x*_1_} between *x*_0_ and *x*_1_ whenever *d*_*X*_ (*x*_0_, *x*_1_) ≤ *ϵ*, a filled-in triangle {*x*_0_, *x*_1_, *x*_2_} if all pairwise distances *d*_*X*_ (*x*_*i*_, *x*_*j*_) are less than or equal to *ϵ*, 0 ≤ *i, j* ≤ 2, and so on for tetrahedra and their higher-dimensional analogs.

Figure 6 panels (a)-(d) provide an example set of points sampled along a circle, and the evolution of its Rips complex as *ϵ* changes. As illustrated in the figure, for small values of *ϵ*, the Rips complex has several disconnected pieces (components) which merge as *ϵ* increases. At particular values of *ϵ*, the Rips complex may have closed loops whose interiors are not yet filled in; these are called holes, or more formally, 1-dimensional homological features [13, Section 2.1]. For instance, the red edge at *ϵ* = 0.71 in Figure 6 panel (b) indicates the creation (the birth) of such a feature. As *ϵ* continues to increase more triangles are added, and holes are filled-in; continuing with the example in Figure 6, the 1-dimensional homological feature born at *ϵ* = 0.71 dies at *ϵ* = 1.88, when the central triangle with a red edge is included in panel (d).

**Figure 6:**
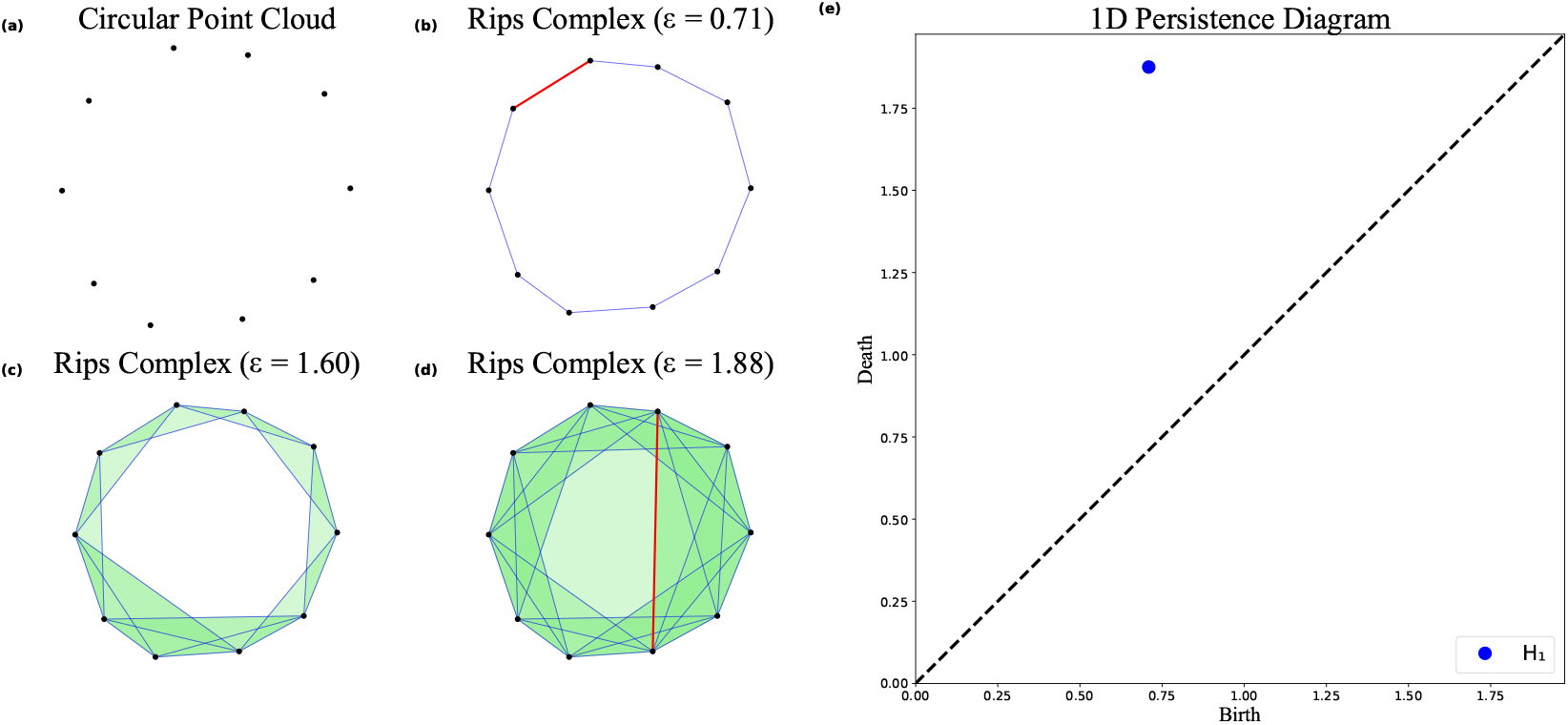
From a point cloud to the 1-dimensional persistence diagram of its Rips filtration. Connected edges in the Rips filtration are drawn in blue, the birth/death of a class is indicated in red, and filled-in triangles are shaded green.

**Figure 7:**
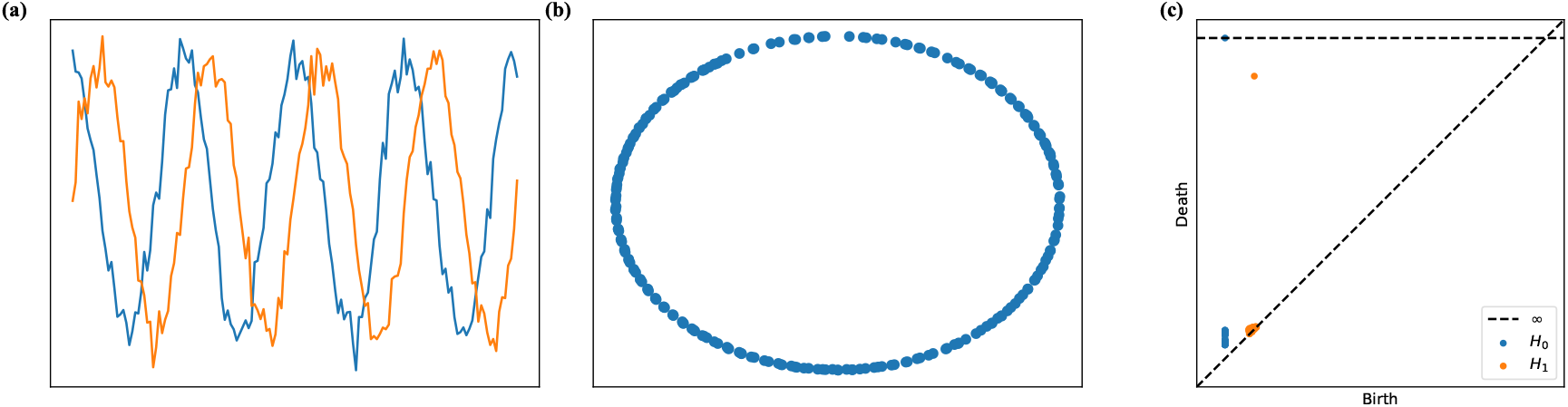
(a) Generated 2-dimensional periodic time series, (b) PCA projection of sliding window point cloud showing circular arrangement, and (c) Computed persistence diagram using *Ripser*.*py* confirming one persistent *H*_1_ feature. Note how one *H*_1_ point rises well above the diagonal, indicating the presences of a “persistent” loop in the data

The persistence diagrams *dgm*_*j*_ for *j* ∈ ℕ, summarize the historical record of *j*-th dimensional homological features *H*_*j*_ in the Rips complex as *ϵ* increases. When *j* = 0 tracking connected components, *j* = 1 tracks holes, *j* = 2 measures voids, and similarly for *j* ≥ 3. More explicitly, *dgm*_*j*_ is a collection of pairs (*b, d*) ∈ ℝ^2^ with *d* ≥ *b* that appear with repetitions; (*b, d*) ∈ *dgm*_*j*_ encodes a *j*-th dimensional homological feature which is born at *ϵ* = *b* and dies at *ϵ* = *d. The persistence* or lifetime of the feature encoded by (*b, d*) is *d* − *b*, and when *dgm*_*j*_ is plotted (Figure 6 panel (e)), points with higher persistence are farther from the diagonal *y* = *x*. In the case of the Rips complex, points in *dgm*_*j*_ with higher persistence indicate the space’s topology from which points were sampled. In contrast, points near the diagonal (low persistence) correspond to gaps and sampling artifacts in the point cloud. Compare Figure 6 panel (e) to Figure 1 panels (d) and (h).

In our implementation, we use `Ripser.py` [35, 4], a highly optimized library for computing persistence diagrams from Rips complexes. A more rigorous treatment of persistent homology can be found in Supplementary Note 3.

### Finding the Optimal Window Size

The two parameters *d* ∈ ℕ and *τ >* 0 used to construct the sliding window point cloud of *F* control the size *dτ* of the window that “slides” along the time series *F* as *t* varies. The window size *dτ*, in turn, influences the shape of the circle described by the sliding window embedding, and hence how effective the persistence diagrams *dgm*_1_ will be in capturing periodic movement (see Supplementary Figure 3). Optimizing these parameters is crucial, and we indicate next the strategies suggested by the theory of sliding windows and persistence [29].

It has been observed that when the window size *dτ* approaches the length of the period of a signal *f*: ℝ −→ ℝ, the roundedness of its sliding window point cloud is maximized. This, in turn, maximizes the persistence of the point farthest from the diagonal in *dgm*_1_, and hence, the strength of the periodicity score computed via persistence diagrams. In particular, if *f* satisfies the identity

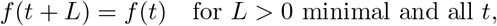

i.e., *f* is *L*-periodic, then its *L*-periodicity is best captured by the persistence diagrams of its sliding window point cloud when:

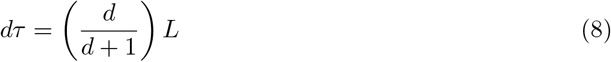

Next we describe how to estimate *L* for a 2-dimensional signal *F* (*t*) = (*f*_*x*_(*t*), *f*_*y*_(*t*)) ∈ ℝ^2^, assuming *F* has been sampled at *N* uniformly spaced time points 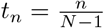. To this end, we define the complex representation *J*[*n*] = *f*_*x*_(*t*_*n*_) + *if*_*y*_(*t*_*n*_) ∈ ℂ of *F*, and use its Discrete Fourier Transform to estimate *L*.

#### Definition 2

(Complex Discrete Fourier Transform). *The Complex Discrete Fourier Transform (CDFT) of the discrete complex-valued signal J is given by*

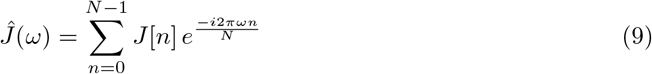

*where ĵ*(*ω*) *is the CDFT coefficient at frequency ω*.

The power spectrum |*ĵ*(*ω*)| of *J* quantifies how strongly each frequency component *ω* is present in the original signal. In particular, let {*ω*_1_, *ω*_2_, …, *ω*_*n*_} be the individual frequency components in the CDFT and let *ω*^∗^ = *argmax*_*ω*_{|*ĵ*(*ω*)|}. Then the period can be estimated as

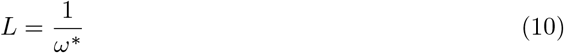

The `numpy.fft` module provides a fast implementation of the Discrete Fourier Transform, and we utilize it for period estimation. The method for estimating the period of trivariate accelerometer time series *F* (*t*) = (*f*_*x*_(*t*), *f*_*y*_(*t*), *f*_*z*_(*t*)) can be found in Supplementary Note 4. Further details of period estimation can also be found in the GitHub repository. With methods for estimating the period, the next step is to find an optimal *d* value.

Traditionally, for non-linear time series analysis, *d* is estimated using false-nearest-neighbors [17], or set to equal the number of prominent peaks (i.e, 70% of the distribution) in the power spectrum [10]. Rather than changing the value from video segment to video segment, we set *d* as a constant value. This has the advantage of comparing point clouds in the same Euclidean Space after taking sliding window embeddings.

In the context of dynamical systems, and according to Takens’ Theorem [34], the embedding dimension *d* must satisfy *d >* 2*m*, where *m* is the dimension of the underlying attractor (e.g., *m* = 1 for a purely periodic signal that concentrates on a topological circle). However, real-world signals are rarely periodic in practice; they often exhibit approximate periodicity or multiple independent oscillatory modes. Using the number of prominent peaks in the power spectrum of the Fourier transform provides a helpful method for modeling the complexity of the underlying dynamics [29, Proposition 5.2]. If *d* is too small, the reconstruction may collapse informative geometric features. To avoid this, we want *d* to exceed twice the number of prominent peaks across all 4-second video segments, ensuring the embedding space has sufficient degrees of freedom to unfold the attractor faithfully. Given the dataset’s sampling rate in this paper, the expected number of prominent peaks in each 4-second window is less than or equal to 10. Therefore, we can conservatively fix *d* = 23 for all analyses.

From a more intuitive and homology-aware perspective, comparing point clouds in different dimensions could lead to altered persistence computations, thus affecting the periodicity scores. We illustrate this phenomenon in Supplementary Fig. 3, using the time series shown in Figure 7, and compute its sliding window embedding across various choices of *d*. We observe how the lifetime of the most persistent *H*_1_ feature changes across multiple ambient dimensions.

Both of these analyses informed our decision to fix the dimension *d*. With our method for estimating the period (*L*) and fixing *d* = 23, we compute *τ* = ^*L*^. With these choices of *d* and *τ*, Figure 7 illustrates the transformation of a 2-dimensional time series into a high-dimensional sliding window point cloud (visualized via PCA), from which Rips persistence diagrams are computed.

### Computing Periodicity Scores

We define *mp*_*n*_(*dgm*_1_) = |*d*_*n*_ − *b*_*n*_| to be the lifetime of the *n*-th most persistent *H*_1_ point in the persistence diagram *dgm*_1_. Lifetime measures the vertical displacement of a point from the diagonal. As shown in Figure 7, we expect only one *H*_1_ feature with high persistence when a time series is periodic. This informs our periodicity score:

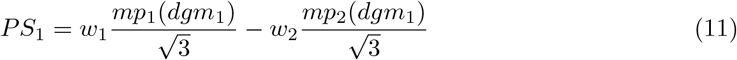

with a weighting function

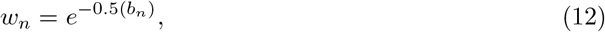

where *b*_*n*_ denotes the birth scale of the *n*-th most persistent *H*_1_ point. A high birth scale in the sliding window point cloud typically indicates that the loop structure, if present, is only detected at a relatively large scale. This often reflects a significant gap or sparsity in the embedding, such that a circular trajectory is not well-sampled or fails to form a tight loop. While the signal may exhibit recurrent behavior, the point cloud lacks sufficient density to generate a loop at an earlier scale, suggesting weak or incomplete periodicity.

This periodicity score measures the offset between the first and second most persistent *H*_1_ features and scales to a value between [0, 1]. This bounded range is a direct consequence of normalizing the sliding window point cloud. The larger this offset is, the more circular the sliding window point cloud is expected to be, as seen in Figure 7. Summarizing periodicity in a single value makes the analysis straightforward and interpretable. In addition to this score representation, we also provide results with an alternative periodicity score that is represented as a vector of the 10 most persistent *H*_1_ features in the persistence diagram:

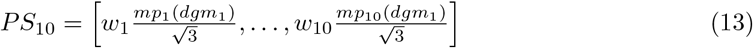

This offers a more robust way of ensuring we see exactly one persistent *H*_1_ point, followed by very low persistent points. The increase in information yields better classification results, with the primary trade-off being a decline in interpretability. The *PS*_1_ representation also minimizes memory usage, which can be particularly valuable when working with large datasets.

### Model Training and Class Imbalance

To reduce the impact of class imbalance, for each video, a random subsample of 4-second intervals without SMMs is analyzed, where the sample taken is equal to *min*(3#(SMM), #(No SMM)) per session. A model trained on an imbalanced dataset can perform poorly on the minority class, limiting generalization. Even when we subsample the intervals with no stereotypy, the imbalance between different SMM classes remains high. To reduce class imbalance on the training dataset, we randomly over-sample the minority classes using *imbalanced-learn*’s `RandomOverSampler`. We used two schemes, wherein both involve duplicating with replacement existing samples from the minority classes until the desired proportions are reached:

- **Uniform Oversampling**: We randomly oversampled the three minority classes, where each class comprised exactly 25% of the total dataset. This oversampling manner was beneficial when training classification models on all four classes, ensuring each class has an equal weight in training.
- **Partial Oversampling**: We over-sampled the minority classes such that the majority class contributes 50% of the samples, while the three minority classes each contribute approximately 16.7%. This scheme is primarily used for binary classification.

### Random Forests

Random Forests are an ensemble classification method that combines predictions of multiple decision tree classifiers. Each decision tree is trained independently on a bootstrap sample drawn from the original training set. Bootstrapping involves randomly sampling the training data with replacement, ensuring each tree has a unique dataset, thereby promoting diversity among the decision trees. Each decision tree learns simple decision rules inferred from data features that can be represented as a search tree, and the random forest makes the majority decision. These models are popular because they are easy to use, scale well to large datasets, handle noise effectively, and are explainable.

### Model Optimization

Random Forest classifiers have numerous hyperparameters that need to be tuned, which can be challenging. We employed Bayesian Optimization to select the best combination of features and find optimal hyperparameters for the Random Forest, thereby preventing overfitting and improving generalization. Rather than performing an exhaustive grid search across all different combinations within the search space, which is computationally expensive, Bayesian Optimization builds a probabilistic surrogate model to guide the search towards more promising regions, significantly reducing the number of evaluations required to identify near-optimal hyperparameters [9]. To properly use Bayesian Optimization, we extracted data and partitioned it into train/validation/test sets, where the training set is used to explore different features and hyperparameters, and the validation set is used to evaluate how well the model fits to unseen data. The testing set is used at the end when the optimal parameters have been chosen. We use the `RandomForestClassifier` from *scikit-learn* for classification, and *scikit-optimize* for hyperparameter tuning via Bayesian optimization. More on Bayesian optimization can be found in Supplementary Note 5.

## Ethics Statement

The study that generated the data featured in the paper was reviewed and approved by IRB Committees at the University of Rhode Island and the Groden Center.

## Data Availability

The data and code used to replicate results is available in the GithHub: https://github.com/ambaye15/AQSM_SW1PerS.git

## Acknowledgments

The National Science Foundation partially supported Austin A. MBaye and Jose A. Perea through CAREER award # DMS-2415445. Autism Speaks, the Nancy Lurie Marks Family Foundation, and grant # R01LM014191 from the National Institutes of Health supported Matthew S. Goodwin. The accelerometry sensors were developed through NSF grant# 0312065.

## Author Contributions

Conceived and designed the experiments: MG, AM, JP, CT; Designed analytical approach: JP, CT, AM; Analyzed the data: All authors; Contributed analysis tools: AM, JP, CT; Wrote the code: AM, CT, JP; Wrote the paper: All authors

