## Supplementary Information for "Automated Quantification of Stereotypical Motor Movements in Autism Using Persistent Homology"

#### Contents

**Supplementary Note 1. Data Information** **2**

**Supplementary Note 2. YOLOv5 Training** **3**

**Supplementary Note 3. Persistent Homology** **3**

**Supplementary Figure 3. Effect of Sliding Window Dimension $d$ on Persistent Homology** **6**

**Supplementary Note 4. Accelerometer Data** **6**

**Supplementary Note 5. Bayesian Optimization** **7**

**Supplementary Note 6. Computational Resources** **8**

**Supplementary Table 2. Experiment 1 - Stratified Set Classification** **9**

**Supplementary Table 3. Experiment 2 - Leave-One-Session-Out** **11**

**Supplementary Table 4. Experiment 3 - Leave-One-Individual-Out** **13**

**Experiment Figures** **13**

**Supplementary Note 7. Application of Methods** **22**

†[2]Department of Mathematics and Khoury College of Computer Sciences, Northeastern University, Boston, MA, USA

‡[3]Department of Mathematics and Computer Sciences, Ursinus College, Collegeville, PA, USA

§[4]Department of Health Sciences, Northeastern University, Boston, MA, USA

### Supplementary Note 1. Data Information

**Supplementary Table. 1** — Overall SMM statistics in our dataset calculated using manual video annotation.

| Participant | Sessions | | No. of Bouts | | | Duration of Bouts (min) | | |
|---|---|---|---|---|---|---|---|---|
| | No. | Dur. (min) | Rock | Flap | Flaprock | Rock | Flap | Flaprock |
| **STUDY 1** | | | | | | | | |
| 1 | 2 | 50.2 | 52 | 2 | 57 | 5.2 | 0.2 | 4.9 |
| 2 | 2 | 32.1 | 1 | 209 | 0 | 0.1 | 8.1 | 0 |
| 3 | 2 | 64.5 | 1 | 52 | 28 | 0.1 | 2.9 | 2.4 |
| 4 | 2 | 38.9 | 55 | 70 | 0 | 12.7 | 6.2 | 0 |
| 5 | 2 | 44.7 | 12 | 75 | 17 | 1.2 | 8.6 | 1.6 |
| 6 | 2 | 55.8 | 112 | 0 | 9 | 31.3 | 0 | 1.6 |
| **STUDY 2** | | | | | | | | |
| 1 | 3 | 49.8 | 77 | 57 | 64 | 7.4 | 6.5 | 5 |
| 2 | 2 | 51.1 | 77 | 33 | 1 | 7.7 | 1 | 0.1 |
| 3 | 2 | 75.9 | 1 | 18 | 0 | 0.02 | 0.5 | 0 |
| 4 | 3 | 87.5 | 46 | 38 | 0 | 13.2 | 2 | 0 |
| 5 | 2 | 55.2 | 68 | 11 | 6 | 10.9 | 1.3 | 0.3 |
| 6 | 1 | 25.3 | 64 | 0 | 6 | 20.2 | 0 | 6 |

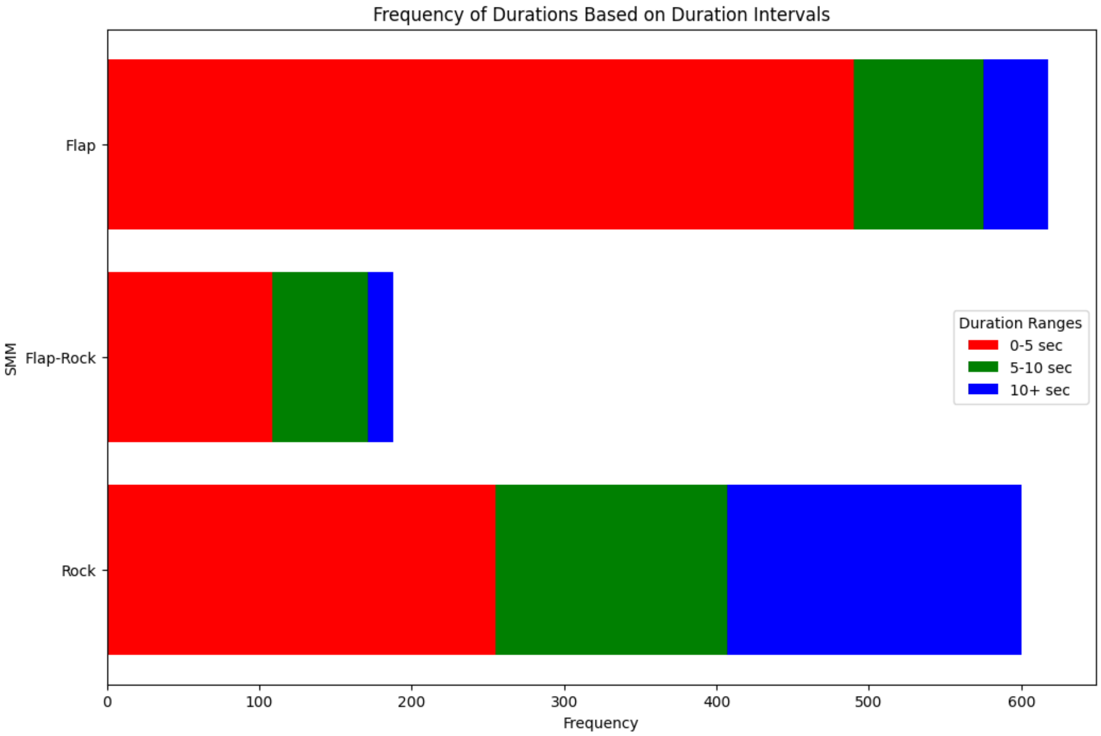

**Supplementary Fig. 1** — Displays the distribution of SMM durations, grouped into time bins. The majority of flapping and flap-rock behaviors occur in short bursts lasting 0–5 seconds, whereas rocking behaviors are more evenly distributed across the 0–5, 5–10, and 10+ second intervals. This suggests for this dataset that rocking tends to manifest in more sustained episodes compared to flapping-related movements, which are typically brief and transient.

### Supplementary Note 2. YOLOv5 Training

We trained a YOLOv5l model via transfer learning. In other words, a pre-trained model of YOLOv5 trained on the COCO [1] dataset is adapted to our custom dataset while still leveraging the model's prior knowledge allowing for a more efficient learning process. A carefully chosen 2,500 video frames were manually annotated by drawing bounding boxes around the person of interest, ensuring we saw the people in an even distribution of situations and poses so tracking would be as accurate as possible. We used an 80/10/10 (train/test/validation) split. Cropping, blurring, and vertical flipping augmentations were applied to the training data to simulate new training examples from which the model could learn. Below shows the results of training the model:

### Supplementary Note 3. Persistent Homology

The goal of persistent homology is to determine the underlying shape $\mathbb{X}$ given a finite, discrete sample $X$. If a time series is periodic, then its sliding window point cloud $X$ is homotopy equivalent to $S^1$ (circular) [2]. The primary method to analyze these high-dimensional point clouds is via persistent homology. However, to understand homology, one needs to understand

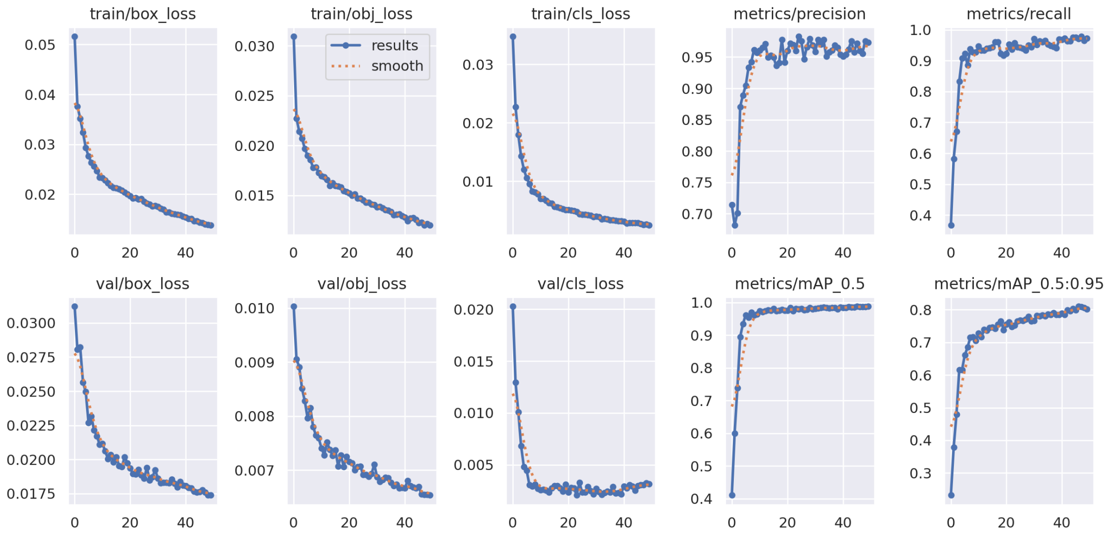

**Supplementary Fig. 2** — YOLOv5l Model Performance Results.

what a simplicial complex is:

**Definition 1** (Simplicial Complex). *A simplicial complex is a collection $K$ of nonempty sets such that:*

- *if* $\sigma \in K$*, then* $0 < \#(\sigma) < \infty$
- *if* $\sigma \in K$ *and* $\varnothing \neq \tilde{\sigma} \subset \sigma$*, then* $\tilde{\sigma} \in K$

*We call* $\sigma$ *an* $n$*-simplex and denote* $K^{(n)}$ *the collection of* $n$*-simplices.*

A 0-simplex represents a vertex; a 1-simplex represents an edge that connects two vertices; a 2-simplex represents a triangle, and analogous interpretations for higher dimensions. Homology, a concept from algebraic topology, is then used to describe the properties of a simplicial complex. These properties include the number of connected components (intuitively, distinct clusters or groups of connected masses), 1-dimensional holes (closed loops bounding empty space), 2-dimensional voids (enclosed shells or surfaces), and higher-dimensional analogues. Let n-chains, denoted as $C_n(K;\mathbb{F})$, be the field $\mathbb{F}$ vector space generated by the n-simplices of a simplicial complex $K$. That is

$$C_n(K;\mathbb{F}) = span_{\mathbb{F}}(K^{(n)})$$

The $n$-chains form a chain complex

$$C_n \xrightarrow{\delta_{n-1}} C_{n-1} \rightarrow \cdots \xrightarrow{\delta_1} C_1 \xrightarrow{\delta_0} C_0$$

where $\delta_i$ is the boundary map given by

$$\delta_i(\sigma) = \sum_{i=0}^{n} (-1)^i (\sigma / \{x_i\}) \in C_{n-1}$$

for $\sigma \in C_n(K)$. The boundary map satisfies $\delta_n \circ \delta_{n+1} = 0 \; \forall n \in \mathbb{N}$.

**Definition 2** (Simplicial Homology Group). *The $n^{th}$ homology group of $K$ is denoted as*

$$H_n(K) = \frac{\ker \delta_n}{\operatorname{im} \delta_{n+1}}$$

The homology of a topological space $X$ consists of a sequence of these vector spaces $H_n(X)$. The rank of $H_n(X)$ counts the number of $n$-dimensional holes. For example, $dim(H_0(X))$ counts the number of connected components, and $dim(H_1(X))$ counts the number of loops.

In practice, we can construct a simplicial complex, known as a *Vietoris-Rips complex*, to compute features on sets of discrete data points.

**Definition 3.** *Given a metric space $(X, \mathbf{d}_X)$, the Vietoris-Rips complex (Rips complex) is defined as:*

$$R_\alpha(X) = \{\, \sigma \subset X \mid 0 < \#(\sigma) < \infty \ , \mathsf{diam}(\sigma) \leq \alpha \,\}$$

*where $\#(\sigma)$ is the cardinality of $\sigma$ and $diam(\sigma)$ is the diameter of $\sigma$.*

Persistent homology tracks the evolution of topological features of complexes such as $R_\alpha(X)$, as the scale parameter $\alpha$ increases. More formally, this is called a *Rips filtration*

**Definition 4.** *Given a metric space $(X, \mathbf{d}_X)$, the Rips Filtration is defined as:*

$$\mathcal{R}(X) = \left\{\, i_{\alpha',\alpha} : R_\alpha(X) \hookrightarrow R_{\alpha'}(X) \,\right\}_{0<\alpha\leq\alpha'}$$

Thus, we obtain a set of sub-complexes of our Rips complex

$$\emptyset \subseteq R_{\alpha_0}(X) \subseteq R_{\alpha_1}(X) \subseteq \cdots \subseteq R_{\alpha_n}(X)$$

Applying homology to each sub-complex in the filtration induces linear maps between vector spaces in each dimension $n$:

$$H_n(R_{\alpha_0}(X)) \to H_n(R_{\alpha_1}(X)) \to \cdots \to H_n(R_{\alpha_n}(X))$$

This structure is known as persistent homology $PH_n(X)$, which tracks how n-dimensional features appear and disappear across distance scales. Each topological feature corresponds to an interval $[b_i, d_i)$, representing the distance scale at which the topological feature is created and when it disappears. The multi-set of such intervals is the barcode or persistence diagram, providing a summary of the dataset's multiscale topological structure.

### Supplementary Figure 3. Effect of Sliding Window Point Cloud Dimension $d$ on Persistent Homology

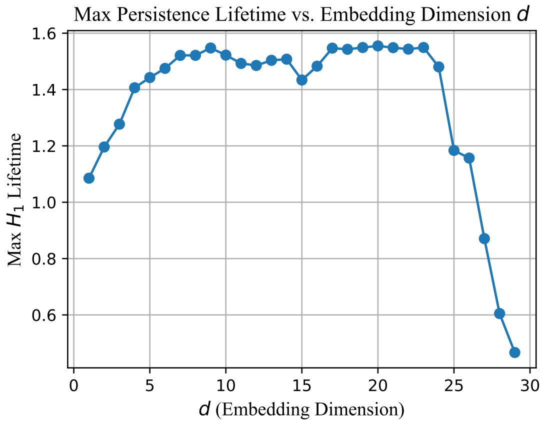

**Supplementary Fig. 3** — Max $H_1$ lifetime of $S\mathbb{W}$ embedding of time series shown in Figure 7 of the manuscript across different ambient dimensions.

### Supplementary Note 4. Accelerometer Data

In order to directly compare results from prior studies and test our *ATSM-SW1PerS* pipeline on a dataset with more temporal resolution, we used the same available data as the videos [3]. Aside from the video data, during both studies, participants wore three 3-axis accelerometers on each wrist and on the torso. Different accelerometers were used in each study. Study 1 used MITes 3-axis sensors recording $\pm 2$ g data at 60 Hz. Study 2 used Wockets 3-axis sensors recording $\pm 4$ g data at 90 Hz. Recordings were done across multiple sessions within each study. These readings serve as a high-resolution signal to test and benchmark our data analysis pipeline. To ensure consistency, each sensor was resampled to 95Hz, and the remainder of the pipeline, from detrending each signal component to computing the periodicity, mostly stays the same with the exception of the period estimation.

#### Estimating Period

In our main manuscript, we estimate the period of a 2-dimensional time series via the Complex Discrete Fourier Transform. However, this does not completely work for the 3-dimensional accelerometer time series. To accurately estimate the period of the entire system, we referred to a recently proposed framework: Learned Accurate Periodic Dictionaries (LAPIS), to estimate the period of multivariate time series [4]. Their method uses a Ramanujan Periodic Dictionary, which consists of structured, periodic basis functions with integer-valued periods. This dictionary provides a sparse and interpretable way to model periodicity, especially in noisy or incomplete signals that are more than 2-dimensional. A complete explanation of methods can be found in their paper and our implementation on GitHub, which offers a robust, and interpretable way to estimate the period of 3-dimensional accelerometer time series $F(t) = (f_x(t), f_y(t), f_z(t))$.

### Supplementary Note 5. Bayesian Optimization

Bayesian Optimization was employed as a strategy to optimize expensive-to-evaluate functions such as optimal feature selection and hyperparameter tuning. Rather than doing a exhaustive grid search across all different combinations within the search space, Bayesian Optimization builds a probabilistic surrogate model to guide the search towards more promising regions.

##### Surrogate Model

The surrogate model, typically a Gaussian Process (GP), provides a probabilistic approximation of the objective function $f(\theta)$, where $\theta$ represents the features and hyper-parameters (the different Mediapipe landmarks and model hyperparameters). A Gaussian Process is defined as:

$$f(\theta) \sim \mathcal{GP}\left(m(\theta), k(\theta, \theta')\right),$$

where:

- $m(\theta)$ is the mean function, often assumed to be zero,
- $k(\theta, \theta')$ is the kernel that encodes the similarity between different parameter settings.

##### Acquisition Function

An acquisition function $a(\theta)$ is used to decide where to sample next, balancing searching uncertain regions of the search space with refining known good areas. One common acquisition function is the Expected Improvement (EI), defined as:

$$\mathrm{EI}(\theta) = \mathbb{E}\left[\max(f(\theta) - f(\theta^+), 0)\right],$$

where $f(\theta^+)$ is the best observed value so far.

##### Optimization Process

The Bayesian Optimization algorithm proceeds as follows:

1. **Initialization:** Evaluate the objective function $f(\theta)$ at a few initial points $\{\theta_i\}_{i=1}^{n}$ to construct the initial surrogate model. We did 30 random starts to build the surrogate model.
2. **Model Update:** Fit or update the Gaussian Process using the current set of observations $\{(\theta_i, f(\theta_i))\}$.
3. **Acquisition Optimization:** Maximize the acquisition function to select the next hyper-parameter setting:
$$\theta_{n+1} = \arg\max_{\theta} \; a(\theta).$$
4. **Evaluation:** Evaluate $f(\theta_{n+1})$ using the actual expensive objective (e.g., F1 score for Random Forest).
5. **Iteration:** Repeat steps 2–4 until a certain number of iterations. We saw after 200 iterations, our evaluations did not improve much, so we set that as our limit.

This approach significantly reduces the number of evaluations required to identify near-optimal hyperparameters, which is particularly beneficial when each evaluation is computationally expensive. For all model training across experiments, we saw using 200 function evaluations and 30 random initializations consistently indicated that retaining all available landmarks led to optimal performance. So when doing any other training, it would make sense to only explore model hypperparametrs, which will reduce the amount of iterations required. One could do 25 random starts and 100 iterations only exploring hyperparametrs drastically reducing optimization time. The parameter space and implementation can be found in our GitHub.

#### Supplementary Note 6. Computational Resources

For fast processing of videos, we utilized Northeastern University's Explorer HPC cluster. For data analysis, processing each video and landmarks can be done in parallel, thus we leveraged SLURM-based job arrays. We split the computations into two parts, video processing and data analysis, each requiring one node. The MediaPipe inference stage used a NVIDIA V100 GPU; the average runtime with these specifications for a single video was approximately 15 minutes. For the data analysis stage, we process 12 landmarks in parallel that require at least 12 CPU cores. To improve efficiency in other computations, we allocated 36 CPU cores and 60GB of memory, with a maximum wall time of 1 hour. On average, each feature extraction would be completed within 5-15 minutes (when using a step size of $1/fps$). When doing a step size of 4 seconds, this time reduces to 1-5 minutes.

#### Supplementary Table 2. Experiment 1 - Stratified Set Classification

**Supplementary Table 2** — Classification performance of Random Forest models using $PS_{10}$ and $PS_1$ periodicity scores across binary and multiclass settings.

| | Pose Estimation | | | | Accelerometer | | | | Accelerometer + Period | | | |
|---|---|---|---|---|---|---|---|---|---|---|---|---|
| **Class** | Precision | Recall | F1 | Support | Precision | Recall | F1 | Support | Precision | Recall | F1 | Support |
| *$PS_{10}$ - Binary Classification* | | | | | | | | | | | | |
| None | 0.83 | 0.95 | 0.88 | 449 | 0.89 | 0.95 | 0.92 | 511 | 0.91 | 0.93 | 0.92 | 511 |
| SMM | 0.91 | 0.72 | 0.80 | 319 | 0.93 | 0.86 | 0.89 | 396 | 0.91 | 0.88 | 0.89 | 396 |
| **Weighted** $F1$ | | | | **0.85** | | | | **0.91** | | | | **0.91** |
| *$PS_1$ - Binary Classification* | | | | | | | | | | | | |
| None | 0.81 | 0.92 | 0.86 | 449 | 0.89 | 0.87 | 0.88 | 511 | 0.91 | 0.89 | 0.90 | 511 |
| SMM | 0.86 | 0.70 | 0.77 | 319 | 0.84 | 0.86 | 0.85 | 396 | 0.86 | 0.88 | 0.87 | 396 |
| **Weighted** $F1$ | | | | **0.82** | | | | **0.87** | | | | **0.89** |
| *$PS_{10}$ - Multiclass Classification* | | | | | | | | | | | | |
| None | 0.80 | 0.92 | 0.86 | 449 | 0.89 | 0.94 | 0.91 | 511 | 0.91 | 0.93 | 0.92 | 511 |
| Rock | 0.87 | 0.78 | 0.82 | 259 | 0.93 | 0.86 | 0.89 | 299 | 0.92 | 0.89 | 0.91 | 299 |
| Flap | 0.22 | 0.09 | 0.13 | 43 | 0.68 | 0.72 | 0.70 | 65 | 0.74 | 0.77 | 0.75 | 65 |
| Flap-Rock | 0.50 | 0.12 | 0.19 | 17 | 0.75 | 0.56 | 0.64 | 32 | 0.76 | 0.59 | 0.67 | 32 |
| **Weighted** $F1$ | | | | **0.79** | | | | **0.88** | | | | **0.89** |
| *$PS_1$ - Multiclass Classification* | | | | | | | | | | | | |
| None | 0.81 | 0.92 | 0.86 | 449 | 0.90 | 0.85 | 0.87 | 511 | 0.91 | 0.88 | 0.89 | 511 |
| Rock | 0.90 | 0.77 | 0.83 | 259 | 0.87 | 0.76 | 0.81 | 299 | 0.90 | 0.85 | 0.88 | 299 |
| Flap | 0.09 | 0.07 | 0.08 | 43 | 0.38 | 0.63 | 0.48 | 65 | 0.57 | 0.78 | 0.66 | 65 |
| Flap-Rock | 0.00 | 0.00 | 0.00 | 17 | 0.22 | 0.34 | 0.27 | 32 | 0.54 | 0.69 | 0.60 | 32 |
| **Weighted** $F1$ | | | | **0.79** | | | | **0.80** | | | | **0.86** |

#### Supplementary Table 3. Experiment 2 - Leave-One-Session-Out

**Supplementary Table 3** — Comparison of our $S\mathbb{W}1PerS$ model trained on accelerometer data (RF-$PS_1$ (Accel) ; RF-$PS_{10}$ (Accel) ), combining frequency-domain features (RF-$PS_1 + Period$ (Accel), RF-$PS_{10} + Period$ (Accel)), and the pose skeleton (RF-$PS_1$ (Pose) ; RF-$PS_{10}$ (Accel)), using a Random Forest classifier against prior work. Performance is evaluated using Accuracy (A) and weighted $F_1$ score, the latter being more informative under class imbalance.

| Study | Method | S1 | | S2 | | S3 | | S4 | | S5 | | S6 | | Mean | |
|---|---|---|---|---|---|---|---|---|---|---|---|---|---|---|---|
| | | $F_1$ | A | $F_1$ | A | $F_1$ | A | $F_1$ | A | $F_1$ | A | $F_1$ | A | $F_1$ | A |
| Study 1 | SVM [3] | 0.73 | 0.86 | 0.36 | 0.85 | 0.50 | 0.94 | 0.73 | 0.67 | 0.44 | 0.75 | 0.46 | 0.87 | 0.54 | 0.82 |
| | RF-RQA [5] | — | 0.71 | — | 0.73 | — | 0.70 | — | 0.92 | — | 0.68 | — | 0.94 | — | 0.78 |
| | CNN-Rad [6] | 0.70 | 0.71 | 0.74 | 0.70 | 0.69 | 0.68 | 0.92 | 0.78 | 0.68 | 0.78 | 0.93 | 0.78 | 0.78 | 0.74 |
| | Frequency-domain CNN [7] | **0.97** | **0.99** | 0.78 | **0.89** | **0.94** | **0.99** | **0.96** | **0.97** | **0.96** | **0.98** | **0.99** | **0.98** | **0.93** | **0.97** |
| | RF-$PS_{10} + Period$ (Accel) | 0.90 | 0.90 | **0.94** | **0.94** | **0.94** | 0.95 | 0.95 | 0.95 | 0.90 | 0.91 | 0.92 | 0.92 | **0.93** | 0.93 |
| | RF-$PS_1 + Period$ (Accel) | 0.90 | 0.90 | 0.91 | 0.91 | **0.94** | 0.94 | 0.92 | 0.92 | 0.87 | 0.88 | 0.93 | 0.93 | 0.91 | 0.91 |
| | RF-$PS_{10}$ (Accel) | 0.89 | 0.89 | **0.94** | **0.94** | **0.94** | 0.94 | 0.95 | 0.95 | 0.90 | 0.91 | 0.90 | 0.90 | 0.92 | 0.92 |
| | RF-$PS_{10}$ (Pose) | 0.78 | 0.81 | 0.68 | 0.75 | 0.68 | 0.76 | 0.77 | 0.78 | 0.70 | 0.77 | 0.88 | 0.87 | 0.75 | 0.79 |
| | RF-$PS_1$ (Accel) | 0.88 | 0.88 | 0.81 | 0.83 | 0.92 | 0.93 | 0.88 | 0.89 | 0.78 | 0.81 | 0.92 | 0.92 | 0.87 | 0.88 |
| | RF-$PS_1$ (Pose) | 0.78 | 0.80 | 0.68 | 0.74 | 0.72 | 0.78 | 0.79 | 0.79 | 0.68 | 0.74 | 0.86 | 0.86 | 0.75 | 0.79 |
| Study 2 | SVM | 0.43 | 0.71 | 0.26 | 0.79 | 0.03 | 0.99 | 0.86 | 0.90 | 0.72 | 0.73 | — | — | 0.46 | 0.82 |
| | RF-RQA | — | 0.80 | — | 0.69 | — | 0.99 | — | 0.95 | — | 0.85 | — | — | — | 0.856 |
| | CNN-Rad | 0.67 | 0.68 | 0.22 | 0.02 | 0.02 | 0.77 | 0.75 | 0.75 | 0.68 | 0.75 | 0.80 | 0.49 | 0.41 | 0.58 |
| | Frequency-domain CNN | **0.96** | **0.97** | **0.95** | **0.98** | **0.85** | **1.00** | **0.98** | **0.99** | **0.94** | **0.97** | — | — | **0.94** | **0.98** |
| | RF-$PS_{10} + Period$ (Accel) | 0.85 | 0.85 | 0.60 | 0.58 | — | — | 0.96 | 0.96 | 0.70 | 0.70 | **0.98** | **0.98** | 0.82 | 0.81 |
| | RF-$PS_1 + Period$ (Accel) | 0.87 | 0.87 | 0.59 | 0.57 | — | — | 0.94 | 0.94 | 0.74 | 0.74 | 0.97 | 0.97 | 0.82 | 0.82 |
| | RF-$PS_{10}$ (Accel) | 0.83 | 0.83 | 0.62 | 0.61 | — | — | 0.93 | 0.93 | 0.71 | 0.71 | **0.98** | **0.98** | 0.81 | 0.81 |
| | RF-$PS_{10}$ (Pose) | 0.83 | 0.84 | 0.71 | 0.75 | — | — | 0.96 | 0.96 | 0.86 | 0.86 | **0.98** | **0.98** | 0.87 | 0.88 |
| | RF-$PS_1$ (Accel) | 0.81 | 0.81 | 0.64 | 0.63 | — | — | 0.92 | 0.91 | 0.64 | 0.64 | **0.98** | **0.98** | 0.80 | 0.79 |
| | RF-$PS_1$ (Pose) | 0.83 | 0.85 | 0.71 | 0.76 | — | — | 0.96 | 0.96 | 0.86 | 0.86 | 0.95 | 0.95 | 0.86 | 0.88 |

### Supplementary Table 4. Experiment 3 - Leave-One-Individual-Out

**Supplementary Table 4** — Binary classification performance (Weighted $F_1$ Scores) of pose trajectories vs. accelerometer signals per participant under Leave-One-Individual-Out evaluation.

| Participant | Pose | | Accelerometer | | | |
|---|---|---|---|---|---|---|
| | $PS_1$ | $PS_{10}$ | $PS_1$ | $PS_{10}$ | $PS_1 + Period$ | $PS_{10} + Period$ |
| 1 | 0.81 | 0.81 | 0.83 | 0.85 | 0.87 | 0.85 |
| 2 | 0.71 | 0.71 | 0.68 | 0.68 | 0.65 | 0.66 |
| 3 | 0.72 | 0.68 | 0.92 | 0.94 | 0.94 | 0.94 |
| 4 | 0.89 | 0.88 | 0.92 | 0.96 | 0.92 | 0.96 |
| 5 | 0.79 | 0.79 | 0.74 | 0.82 | 0.81 | 0.82 |
| 6 | 0.89 | 0.91 | 0.94 | 0.93 | 0.93 | 0.90 |
| **Mean** | **0.80** | **0.80** | **0.84** | **0.86** | **0.85** | **0.86** |

### Experiment Figures

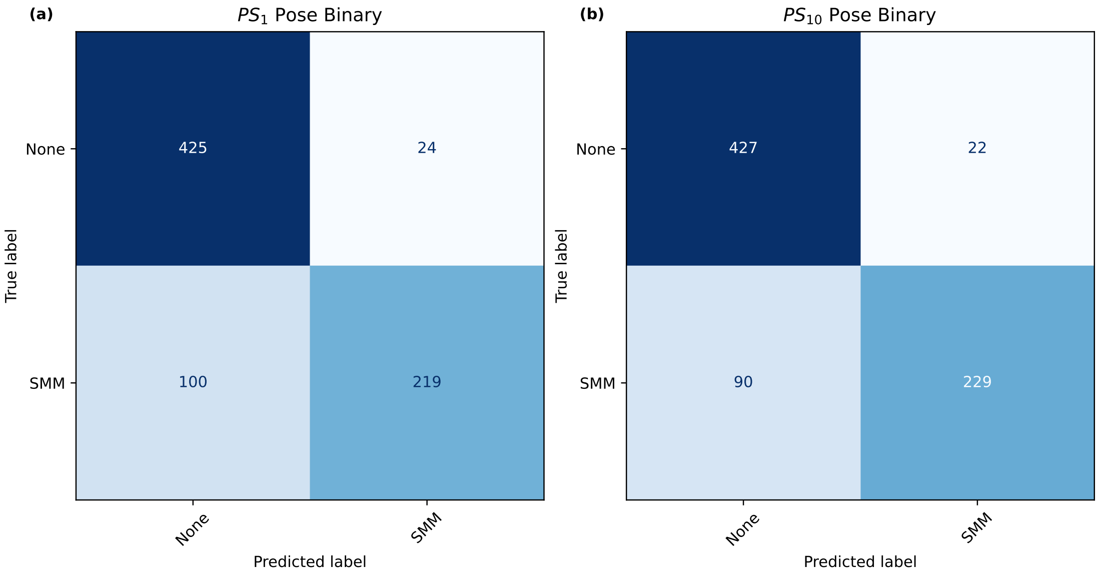

**Supplementary Fig. 4** — Experiment 1 confusion matrices showing binary model's performance using pose-based landmarks. We see both $PS_1$ (a), and $PS_{10}(b)$ have similar performance even though we have an order of magnitude less information using $PS_1$ scores. It is important to note that our model rarely hallucinates periodicity which is important in clinical settings.

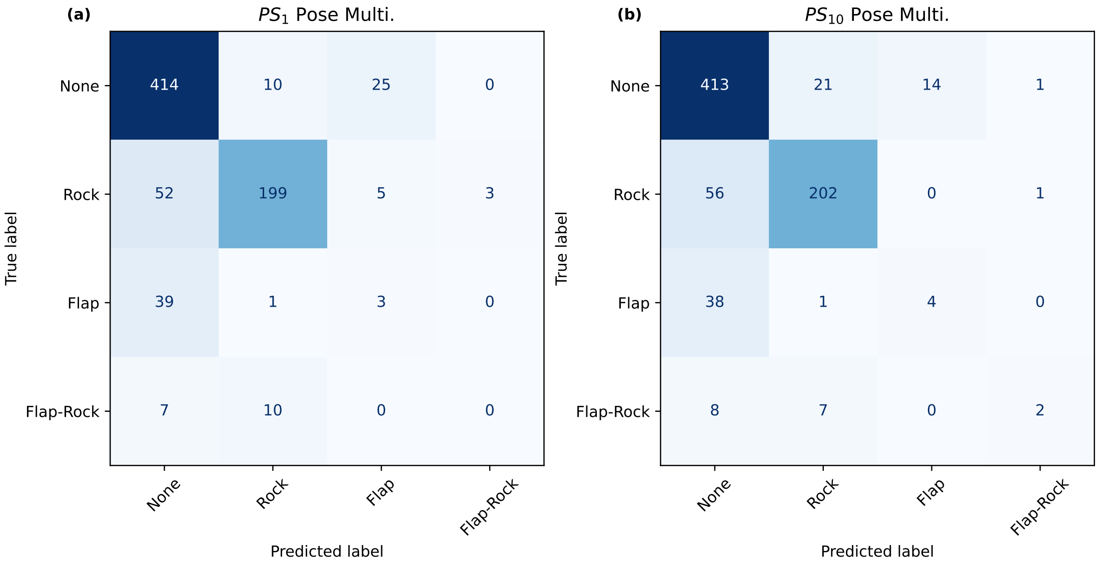

**Supplementary Fig. 5** — Experiment 1 confusion matrices showing multiclass model's performance using pose-based landmarks. Both $PS_1$ (a), and $PS_{10}(b)$ have similar performance. For both panels, we see that near all of flapping instances are misclassified as no stereotypy.

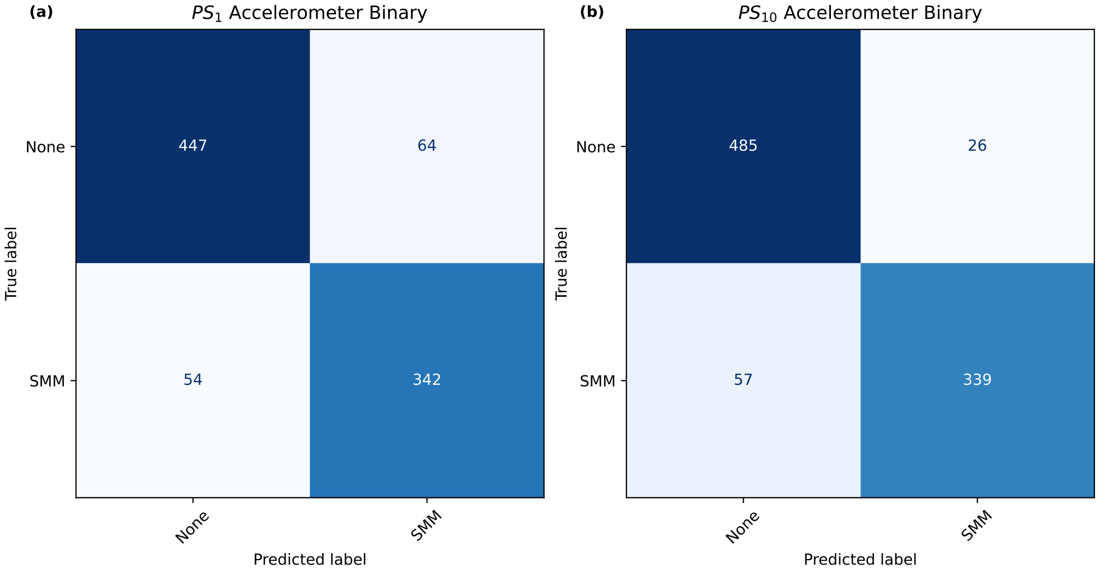

**Supplementary Fig. 6** — Experiment 1 confusion matrices showing binary model's performance using accelerometer-based sensors. We again see both $PS_1$ (a), and $PS_{10}(b)$ have similar performance.

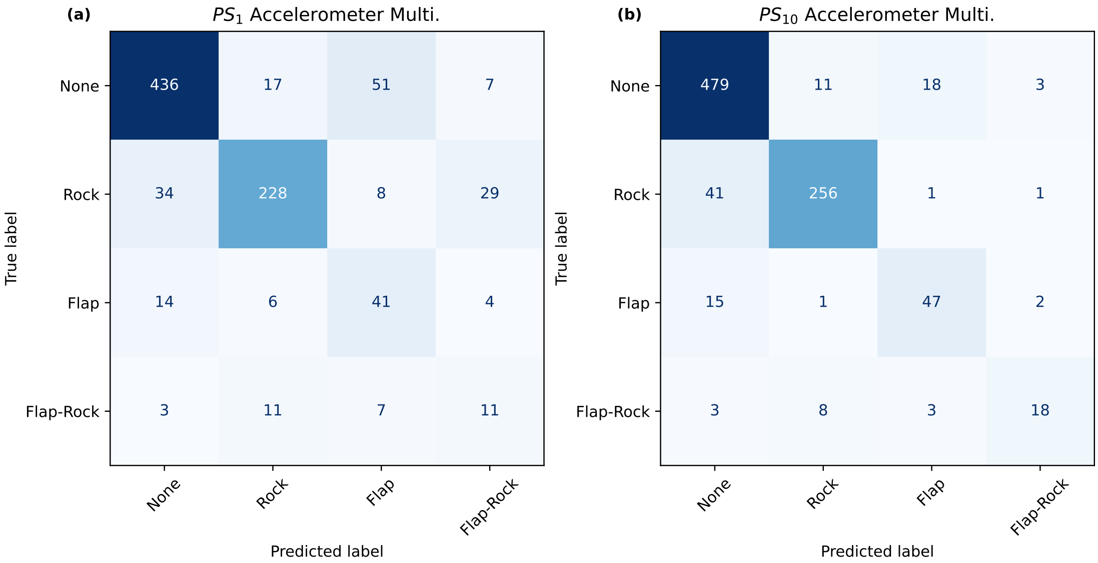

**Supplementary Fig. 7** — Experiment 1 confusion matrices showing multiclass model's performance using accelerometer-based sensors. Both $PS_1$ (a), and $PS_{10}(b)$ have similar performance. For both panels, we see the model's ability to detect repetitive flapping motions is much better than using pose-based landmarks. This is likely due to accelerometers not being affected by occlusions.

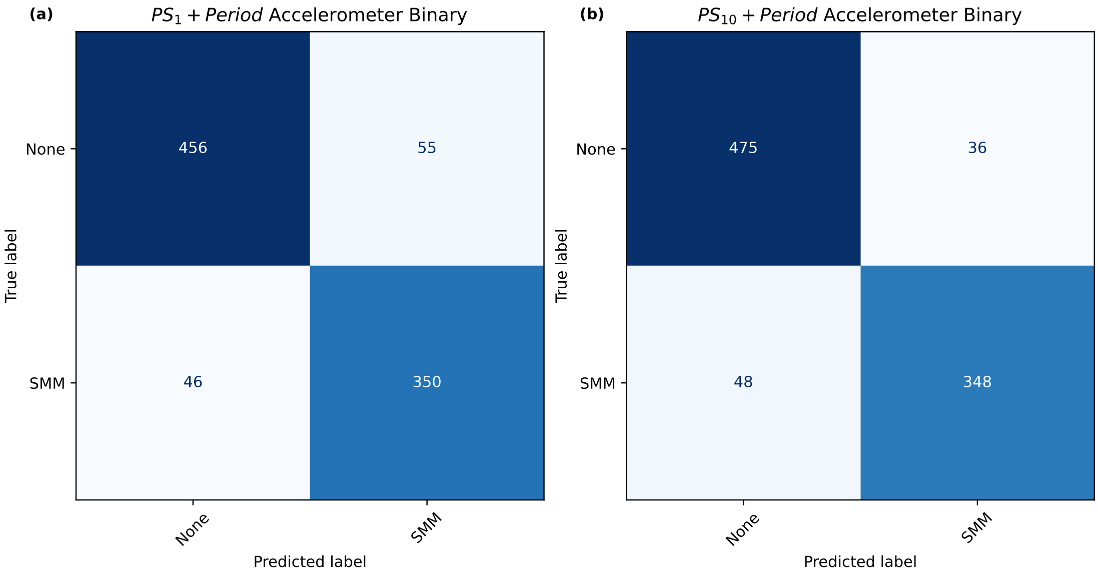

**Supplementary Fig. 8** — Experiment 1 confusion matrices showing binary model's performance using accelerometer-based sensors using frequency-domain and our periodicity scores.

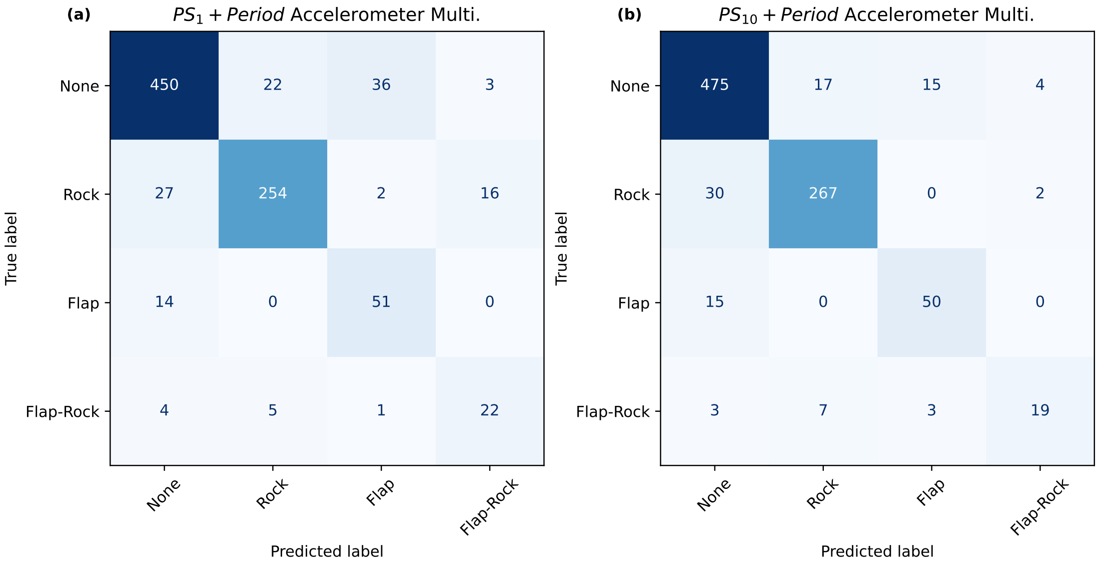

**Supplementary Fig. 9** — Experiment 1 confusion matrices showing multiclass model's performance using accelerometer-based sensors using frequency-domain and our periodicity scores.

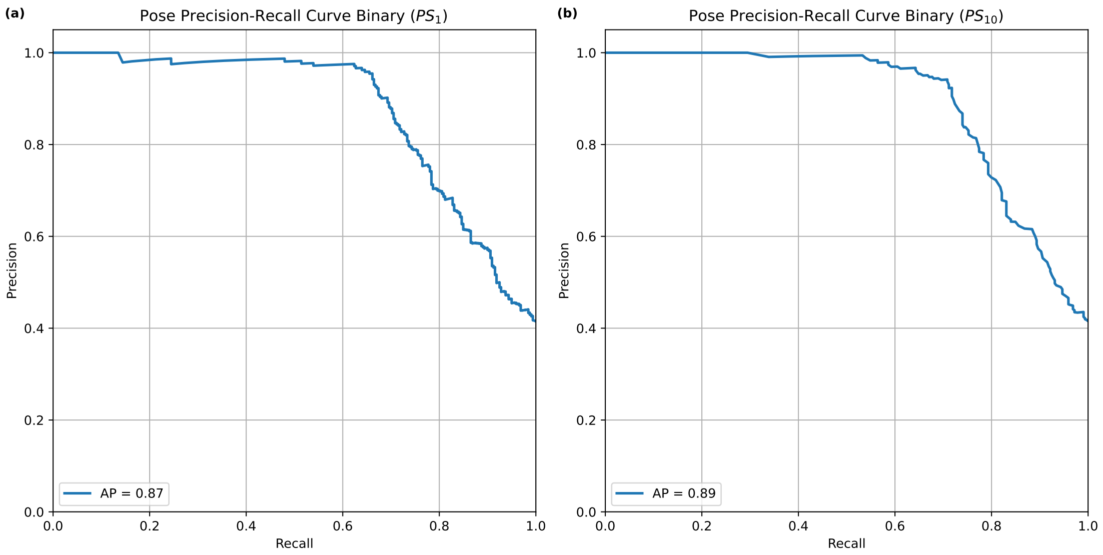

**Supplementary Fig. 10** — Experiment 1 binary classification precision–recall curves for each periodicity score method using pose-based landmarks. Panel a shows results for the $PS_1$ model (AP = 0.87), while panel b displays the $PS_{10}$ model (AP = 0.89). Both curves reflect the trade-off between recall and precision across decision thresholds, with high average precision (AP) indicating strong separability between classes.

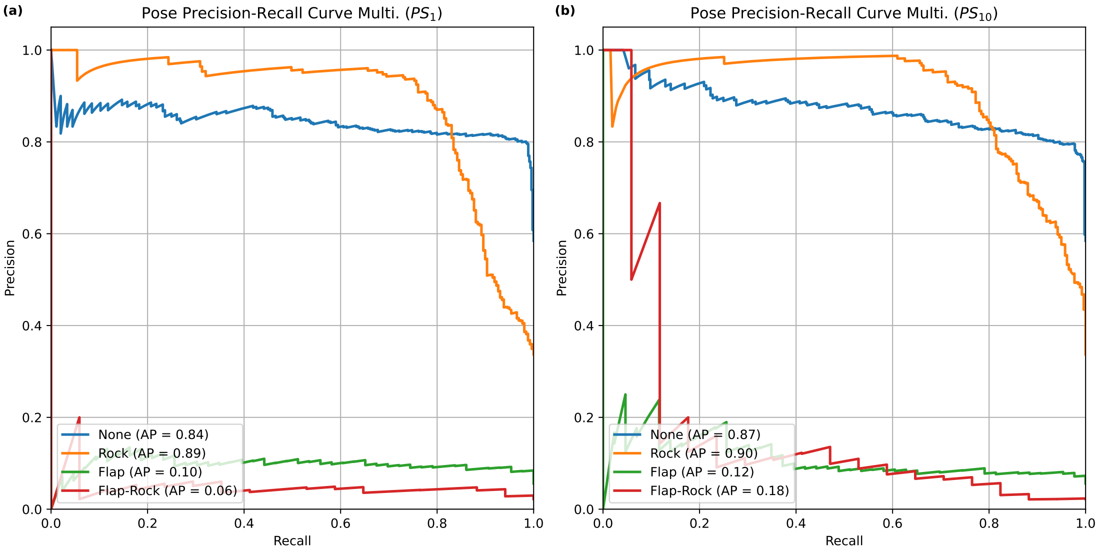

**Supplementary Fig. 11** — Experiment 1 multiclass classification precision–recall curves for each periodicity score method using pose-based landmarks.

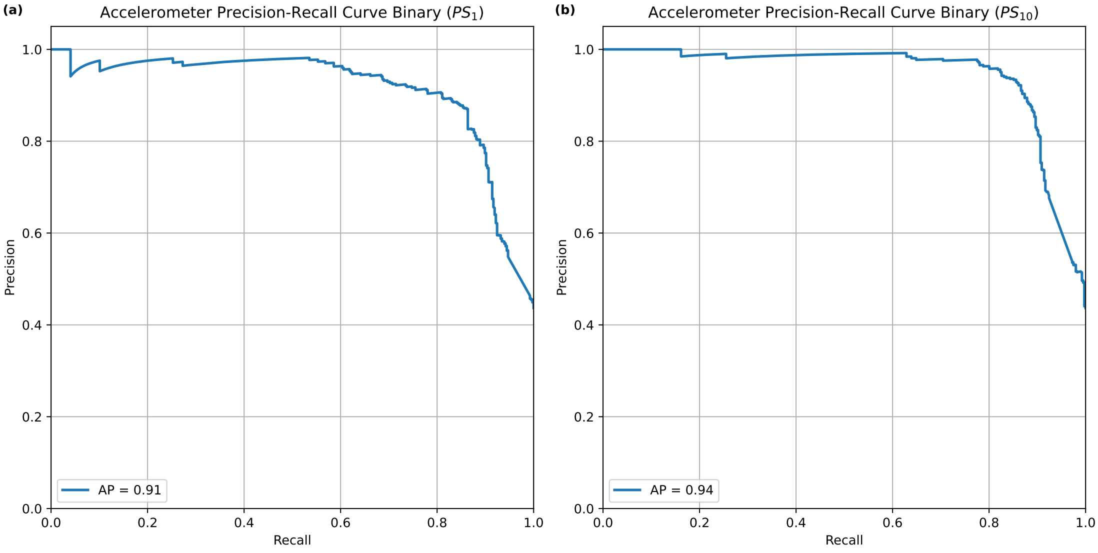

**Supplementary Fig. 12** — Experiment 1 binary classification precision–recall curves for each periodicity score method using accelerometer-based sensors. Panel (a) shows results for the $PS_1$ model (AP = 0.91), while panel (b) displays the $PS_{10}$ model (AP = 0.94). Both curves show that using the accelerometer-based sensors slightly increases performance over the pose-based landmarks.

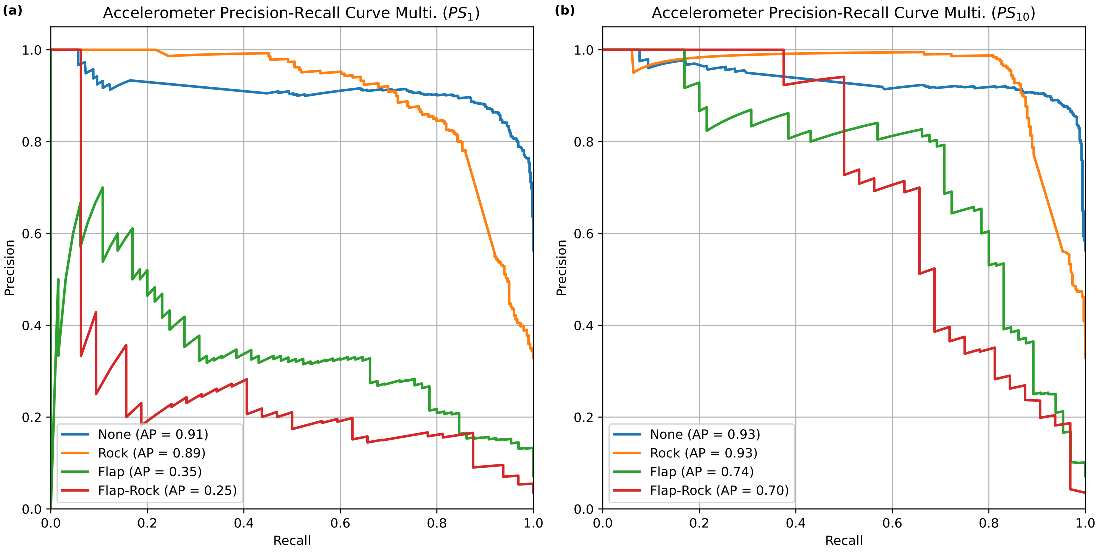

**Supplementary Fig. 13** — Experiment 1 multiclass classification precision–recall curves for each periodicity score method using accelerometer-based sensors.

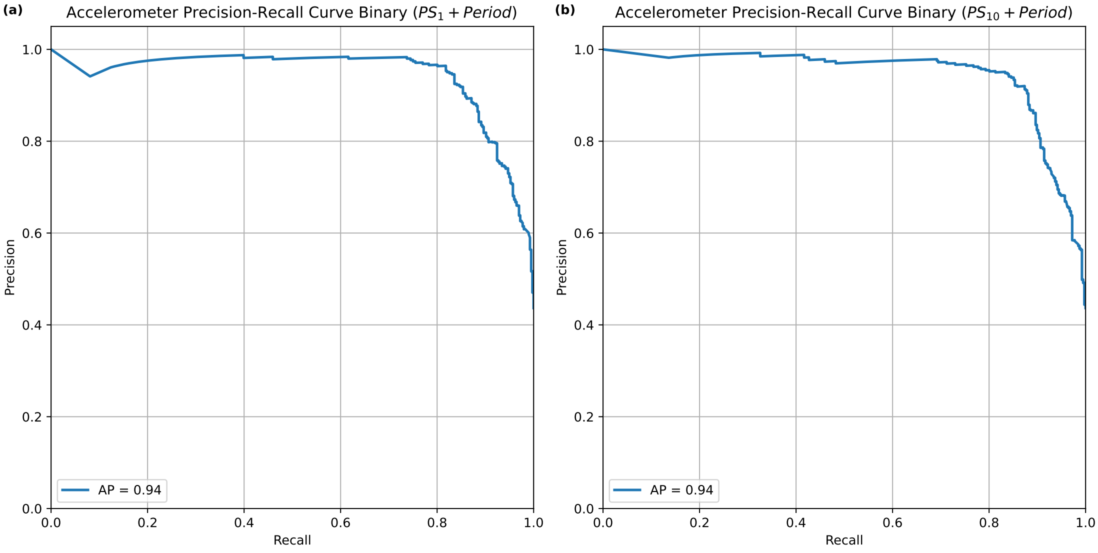

**Supplementary Fig. 14** — Experiment 1 binary classification precision–recall curves for each periodicity score method using accelerometer-based sensors.

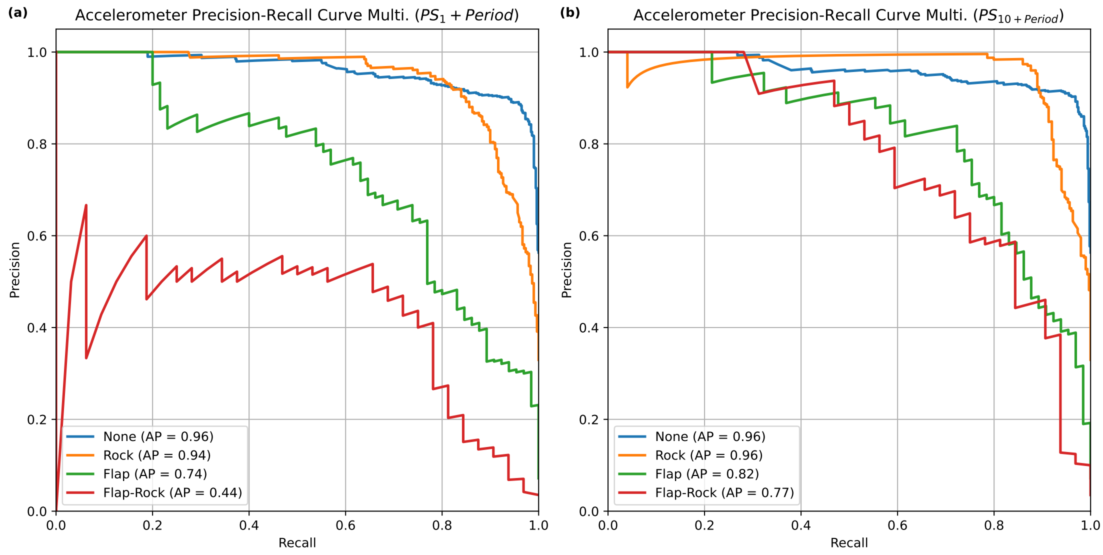

**Supplementary Fig. 15** — Experiment 1 multiclass classification precision–recall curves for each periodicity score method combined with frequency-domain features using accelerometer-based sensors.

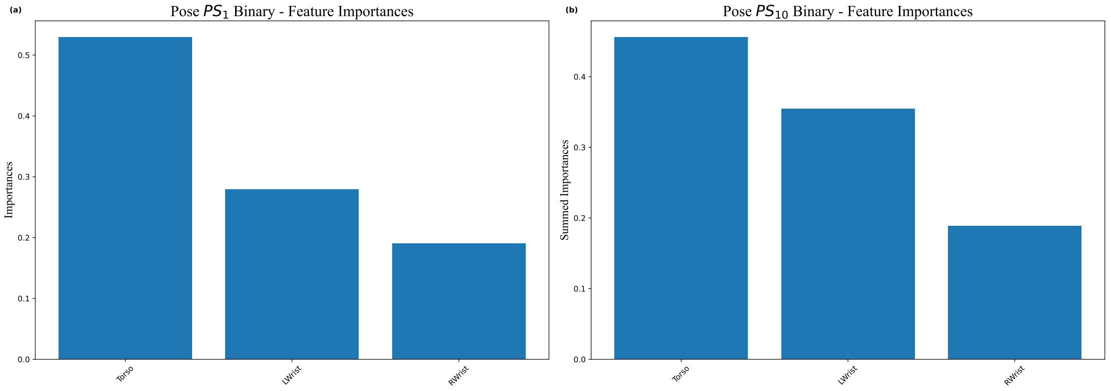

**Supplementary Fig. 16** — Experiment 1 feature importances for binary classification using accelerometer-based sensors. Panel (a) shows Torso contributes the most to the model's decisions. In contrast, panel (b) reveals that all sensors have more equal contribution. This prominence in torso-related features aligns with the pose-based landmark feature importances and prior findings [5].

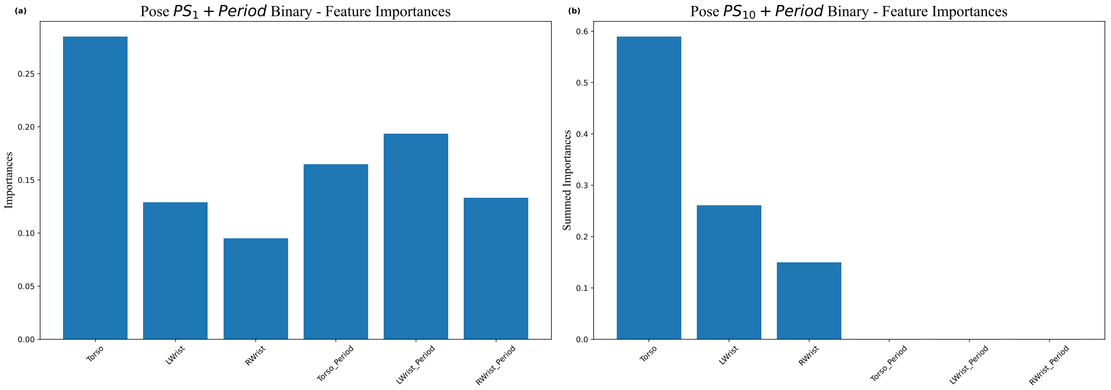

**Supplementary Fig. 17** — Experiment 1 feature importances for binary classification using accelerometer-based sensors using both periodicity scores and period features. Panel (a) shows all features make a contribution to the model's decisions. This time, panel (b) shows that the Torso is primarily responsible in the full-resolution $PS_{10}$ model.

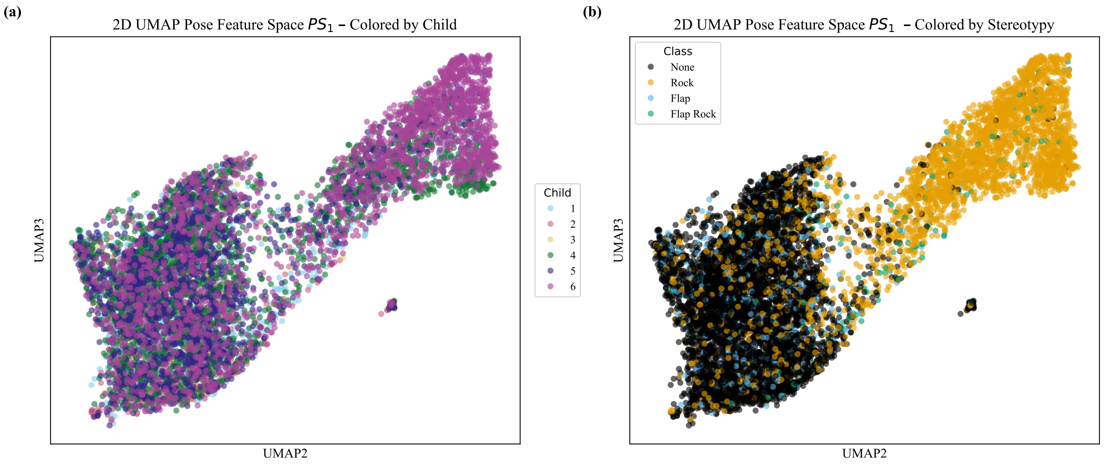

**Supplementary Fig. 18** — Experiment 4 *UMAP* embeddings of pose-based landmark $PS_1$ feature space colored by (a) participant ID and (b) SMM class. Same organization as the rest

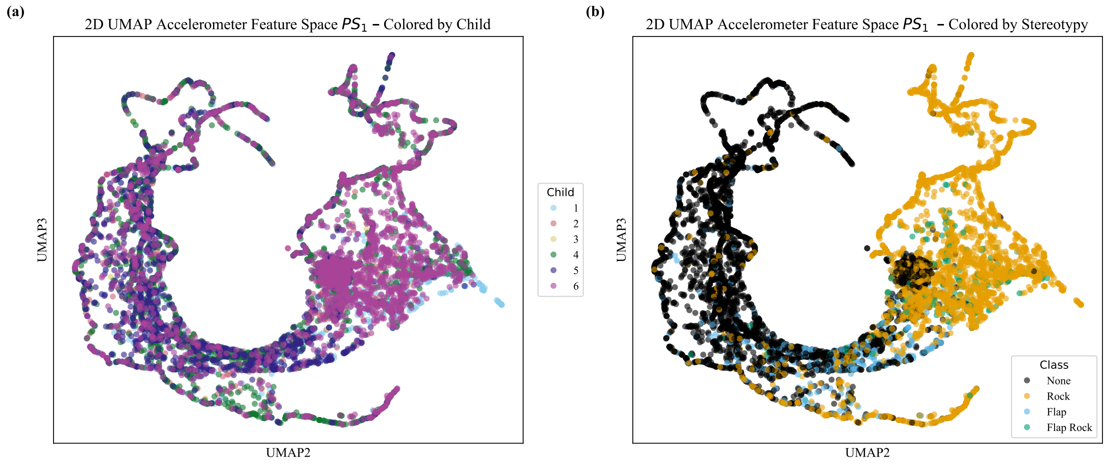

**Supplementary Fig. 19** — Experiment 4 *UMAP* embeddings of accelerometer-based sensor $PS_1$ feature space colored by (a) participant ID and (b) SMM class. Note that the feature spaces are subsampled down for ease of visualization. In (a), the overlap across individuals suggests no variability based on ID. In contrast, (b) shows clear organization based on SMM class, indicating that despite individual variability, our topological features encode behaviorally meaningful patterns that generalize across participants even with accelerometer-based sensors so it is not specific to pose-based landmarks.

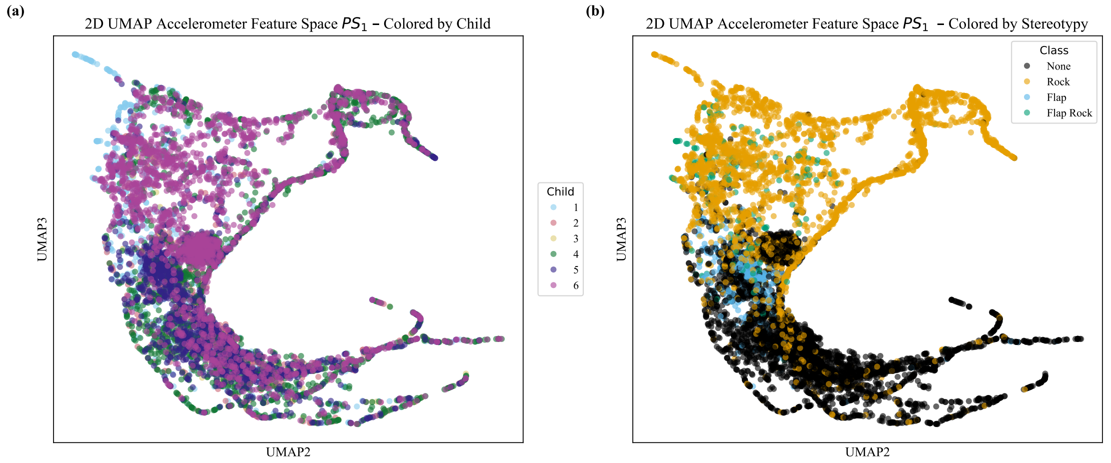

**Supplementary Fig. 20** — Experiment 4 *UMAP* embeddings of accelerometer-based sensor $PS_{10}$ feature space colored by (a) participant ID and (b) SMM class.

### Supplementary Note 7. Application of Methods

#### Illustrative Example: Persistence Changes Within a Participant Over Time

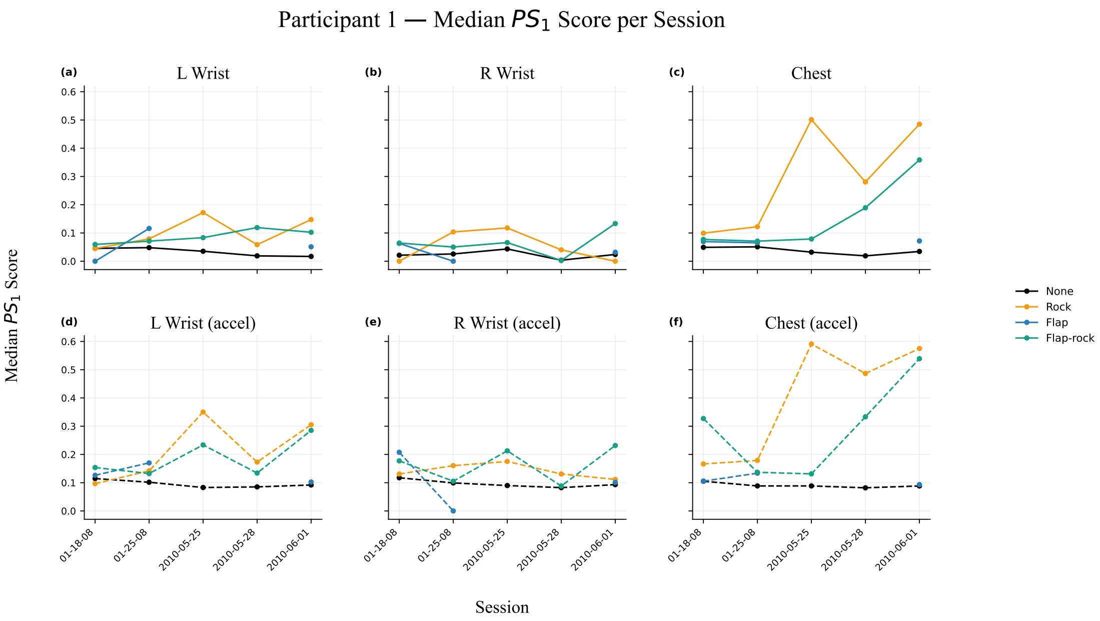

**Supplementary Fig. 21** — Evolution of median $PS_1$ score derived from $S\mathbb{W}1PerS$ pipeline on representative set of pose landmarks.

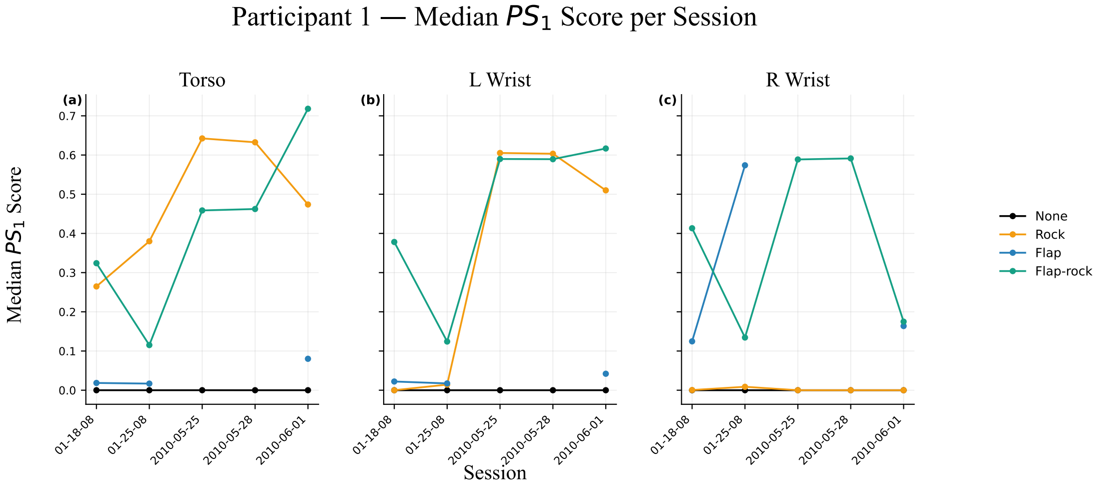

**Supplementary Fig. 22** — Evolution of median $PS_1$ score derived from $S\mathbb{W}1PerS$ pipeline on accelerometer sensors

##### Periodicity Score Summary Across Participants, Sessions, and Annotations

**Supplementary Table 5** — Pose Landmarks Median ± SD $PS_1$ Scores per Sensor, Stratified by Participant, Session, and Annotation Part I

| Participant | Session | Annotation | RWrist | LWrist | Chest | RWrist Accel | LWrist Accel | Chest Accel |
|---|---|---|---|---|---|---|---|---|
| 1 | 01-18-08 | None | $0.021 \pm 0.115$ | $0.045 \pm 0.109$ | $0.049 \pm 0.117$ | $0.117 \pm 0.116$ | $0.115 \pm 0.104$ | $0.105 \pm 0.104$ |
| | | Rock | $0.000 \pm 0.109$ | $0.045 \pm 0.118$ | $0.099 \pm 0.153$ | $0.131 \pm 0.135$ | $0.097 \pm 0.113$ | $0.166 \pm 0.161$ |
| | | Flap | $0.063 \pm 0.085$ | $0.000 \pm 0.001$ | $0.070 \pm 0.060$ | $0.208 \pm 0.058$ | $0.126 \pm 0.024$ | $0.105 \pm 0.081$ |
| | | Flap-Rock | $0.064 \pm 0.099$ | $0.059 \pm 0.122$ | $0.077 \pm 0.119$ | $0.177 \pm 0.150$ | $0.154 \pm 0.146$ | $0.327 \pm 0.204$ |
| | 01-25-08 | None | $0.026 \pm 0.110$ | $0.048 \pm 0.101$ | $0.051 \pm 0.100$ | $0.099 \pm 0.101$ | $0.102 \pm 0.093$ | $0.089 \pm 0.087$ |
| | | Rock | $0.104 \pm 0.179$ | $0.079 \pm 0.151$ | $0.122 \pm 0.170$ | $0.160 \pm 0.180$ | $0.142 \pm 0.109$ | $0.179 \pm 0.143$ |
| | | Flap | $0.000 \pm 0.000$ | $0.116 \pm 0.157$ | $0.065 \pm 0.138$ | $0.000 \pm 0.089$ | $0.170 \pm 0.085$ | $0.134 \pm 0.084$ |
| | | Flap-Rock | $0.050 \pm 0.100$ | $0.071 \pm 0.101$ | $0.071 \pm 0.127$ | $0.105 \pm 0.100$ | $0.132 \pm 0.113$ | $0.137 \pm 0.111$ |
| | 05-25-10 | None | $0.043 \pm 0.104$ | $0.035 \pm 0.111$ | $0.032 \pm 0.101$ | $0.090 \pm 0.090$ | $0.083 \pm 0.084$ | $0.089 \pm 0.084$ |
| | | Rock | $0.118 \pm 0.145$ | $0.172 \pm 0.182$ | $0.501 \pm 0.256$ | $0.175 \pm 0.136$ | $0.350 \pm 0.177$ | $0.591 \pm 0.199$ |
| | | Flap-Rock | $0.066 \pm 0.098$ | $0.083 \pm 0.086$ | $0.079 \pm 0.111$ | $0.213 \pm 0.118$ | $0.234 \pm 0.155$ | $0.131 \pm 0.090$ |
| | 05-28-10 | None | $0.004 \pm 0.098$ | $0.019 \pm 0.089$ | $0.019 \pm 0.088$ | $0.083 \pm 0.117$ | $0.085 \pm 0.107$ | $0.082 \pm 0.090$ |
| | | Rock | $0.040 \pm 0.112$ | $0.059 \pm 0.143$ | $0.281 \pm 0.210$ | $0.131 \pm 0.172$ | $0.173 \pm 0.152$ | $0.487 \pm 0.171$ |
| | | Flap-Rock | $0.003 \pm 0.129$ | $0.119 \pm 0.108$ | $0.189 \pm 0.153$ | $0.088 \pm 0.062$ | $0.134 \pm 0.121$ | $0.333 \pm 0.187$ |
| | 06-01-10 | None | $0.024 \pm 0.117$ | $0.017 \pm 0.103$ | $0.034 \pm 0.097$ | $0.093 \pm 0.115$ | $0.092 \pm 0.095$ | $0.088 \pm 0.089$ |
| | | Rock | $0.000 \pm 0.072$ | $0.147 \pm 0.165$ | $0.485 \pm 0.277$ | $0.112 \pm 0.134$ | $0.305 \pm 0.159$ | $0.575 \pm 0.182$ |
| | | Flap | $0.032 \pm 0.101$ | $0.051 \pm 0.101$ | $0.072 \pm 0.120$ | $0.101 \pm 0.117$ | $0.102 \pm 0.096$ | $0.093 \pm 0.098$ |
| | | Flap-Rock | $0.133 \pm 0.167$ | $0.103 \pm 0.137$ | $0.359 \pm 0.243$ | $0.232 \pm 0.171$ | $0.285 \pm 0.173$ | $0.539 \pm 0.166$ |
| 2 | 01-18-08 | None | $0.048 \pm 0.113$ | $0.010 \pm 0.090$ | $0.022 \pm 0.107$ | $0.105 \pm 0.101$ | $0.115 \pm 0.122$ | $0.100 \pm 0.096$ |
| | | Flap | $0.037 \pm 0.134$ | $0.058 \pm 0.104$ | $0.043 \pm 0.094$ | $0.084 \pm 0.093$ | $0.105 \pm 0.138$ | $0.119 \pm 0.133$ |
| | 01-24-08 | None | $0.054 \pm 0.112$ | $0.028 \pm 0.103$ | $0.053 \pm 0.118$ | $0.092 \pm 0.098$ | $0.109 \pm 0.107$ | $0.092 \pm 0.112$ |
| | | Rock | $0.000 \pm 0.000$ | $0.014 \pm 0.110$ | $0.095 \pm 0.088$ | $0.000 \pm 0.084$ | $0.187 \pm 0.121$ | $0.174 \pm 0.170$ |
| | | Flap | $0.039 \pm 0.121$ | $0.024 \pm 0.088$ | $0.076 \pm 0.130$ | $0.124 \pm 0.113$ | $0.173 \pm 0.118$ | $0.068 \pm 0.060$ |
| | 06-04-10 | None | $0.000 \pm 0.103$ | $0.003 \pm 0.094$ | $0.018 \pm 0.106$ | $0.080 \pm 0.107$ | $0.094 \pm 0.110$ | $0.081 \pm 0.091$ |
| | | Rock | $0.000 \pm 0.153$ | $0.016 \pm 0.167$ | $0.191 \pm 0.256$ | $0.108 \pm 0.156$ | $0.103 \pm 0.159$ | $0.268 \pm 0.218$ |
| | | Flap | $0.000 \pm 0.065$ | $0.148 \pm 0.057$ | $0.016 \pm 0.064$ | $0.042 \pm 0.050$ | $0.088 \pm 0.080$ | $0.042 \pm 0.080$ |
| | | Flap-Rock | $0.000 \pm 0.207$ | $0.000 \pm 0.025$ | $0.050 \pm 0.127$ | $0.068 \pm 0.079$ | $0.059 \pm 0.153$ | $0.153 \pm 0.264$ |
| | 06-02-11 | None | $0.000 \pm 0.099$ | $0.000 \pm 0.092$ | $0.006 \pm 0.110$ | $0.085 \pm 0.136$ | $0.078 \pm 0.125$ | $0.098 \pm 0.104$ |
| | | Rock | $0.000 \pm 0.094$ | $0.000 \pm 0.110$ | $0.000 \pm 0.098$ | $0.068 \pm 0.129$ | $0.097 \pm 0.113$ | $0.073 \pm 0.084$ |
| 3 | 01-18-08 | None | $0.003 \pm 0.096$ | $0.013 \pm 0.114$ | $0.016 \pm 0.120$ | $0.097 \pm 0.108$ | $0.087 \pm 0.101$ | $0.082 \pm 0.091$ |
| | | Flap | $0.033 \pm 0.043$ | $0.023 \pm 0.047$ | $0.018 \pm 0.060$ | $0.127 \pm 0.066$ | $0.099 \pm 0.048$ | $0.075 \pm 0.065$ |
| | | Flap-Rock | $0.060 \pm 0.086$ | $0.067 \pm 0.149$ | $0.064 \pm 0.119$ | $0.165 \pm 0.147$ | $0.214 \pm 0.151$ | $0.213 \pm 0.179$ |
| | 02-08-08 | None | $0.014 \pm 0.106$ | $0.035 \pm 0.098$ | $0.032 \pm 0.082$ | $0.096 \pm 0.091$ | $0.093 \pm 0.098$ | $0.093 \pm 0.088$ |
| | | Flap | $0.048 \pm 0.105$ | $0.057 \pm 0.103$ | $0.062 \pm 0.088$ | $0.171 \pm 0.132$ | $0.161 \pm 0.073$ | $0.068 \pm 0.064$ |
| | | Flap-Rock | $0.026 \pm 0.089$ | $0.037 \pm 0.138$ | $0.038 \pm 0.116$ | $0.116 \pm 0.109$ | $0.097 \pm 0.078$ | $0.149 \pm 0.085$ |

**Supplementary Table 6** — Pose Landmarks Median ± SD $PS_1$ Scores per Sensor, Stratified by Participant, Session, and Annotation Part II

| Participant | Session | Annotation | RWrist | LWrist | Chest | RWrist Accel | LWrist Accel | Chest Accel |
|---|---|---|---|---|---|---|---|---|
| 4 | 01-17-08 | None | 0.039 ± 0.100 | 0.051 ± 0.106 | 0.049 ± 0.112 | 0.114 ± 0.095 | 0.104 ± 0.105 | 0.102 ± 0.109 |
| | | Rock | 0.107 ± 0.144 | 0.150 ± 0.163 | 0.221 ± 0.209 | 0.146 ± 0.114 | 0.194 ± 0.128 | 0.368 ± 0.164 |
| | | Flap | 0.098 ± 0.127 | 0.071 ± 0.124 | 0.084 ± 0.136 | 0.128 ± 0.092 | 0.097 ± 0.089 | 0.089 ± 0.079 |
| | 02-07-08 | None | 0.048 ± 0.102 | 0.033 ± 0.115 | 0.041 ± 0.111 | 0.102 ± 0.086 | 0.099 ± 0.111 | 0.093 ± 0.083 |
| | | Rock | 0.143 ± 0.166 | 0.104 ± 0.144 | 0.320 ± 0.224 | 0.185 ± 0.146 | 0.137 ± 0.122 | 0.325 ± 0.160 |
| | | Flap | 0.055 ± 0.111 | 0.053 ± 0.106 | 0.070 ± 0.114 | 0.102 ± 0.102 | 0.145 ± 0.115 | 0.085 ± 0.093 |
| | 04-27-10 | None | 0.039 ± 0.104 | 0.010 ± 0.087 | 0.050 ± 0.106 | 0.092 ± 0.089 | 0.103 ± 0.103 | 0.089 ± 0.082 |
| | | Rock | 0.468 ± 0.242 | 0.191 ± 0.171 | 0.688 ± 0.224 | 0.362 ± 0.162 | 0.191 ± 0.122 | 0.393 ± 0.170 |
| | | Flap | 0.086 ± 0.069 | 0.121 ± 0.190 | 0.083 ± 0.089 | 0.080 ± 0.051 | 0.051 ± 0.086 | 0.086 ± 0.056 |
| | 05-11-10 | None | 0.022 ± 0.102 | 0.034 ± 0.097 | 0.042 ± 0.106 | 0.094 ± 0.094 | 0.094 ± 0.096 | 0.087 ± 0.091 |
| | | Rock | 0.177 ± 0.182 | 0.116 ± 0.157 | 0.529 ± 0.242 | 0.121 ± 0.109 | 0.198 ± 0.137 | 0.307 ± 0.179 |
| | | Flap | 0.047 ± 0.121 | 0.072 ± 0.086 | 0.086 ± 0.081 | 0.107 ± 0.143 | 0.116 ± 0.085 | 0.097 ± 0.079 |
| 5 | 01-16-08 | None | 0.039 ± 0.112 | 0.042 ± 0.099 | 0.022 ± 0.096 | 0.120 ± 0.113 | 0.106 ± 0.099 | 0.082 ± 0.088 |
| | | Rock | 0.008 ± 0.120 | 0.120 ± 0.191 | 0.119 ± 0.198 | 0.045 ± 0.081 | 0.136 ± 0.124 | 0.055 ± 0.153 |
| | | Flap | 0.012 ± 0.130 | 0.019 ± 0.126 | 0.021 ± 0.122 | 0.088 ± 0.134 | 0.100 ± 0.109 | 0.084 ± 0.088 |
| | | Flap-Rock | 0.000 ± 0.076 | 0.000 ± 0.113 | 0.269 ± 0.216 | 0.108 ± 0.103 | 0.083 ± 0.087 | 0.190 ± 0.201 |
| | 02-08-08 | None | 0.026 ± 0.095 | 0.034 ± 0.115 | 0.042 ± 0.102 | 0.104 ± 0.118 | 0.113 ± 0.113 | 0.098 ± 0.094 |
| | | Flap | 0.000 ± 0.098 | 0.050 ± 0.105 | 0.039 ± 0.101 | 0.119 ± 0.131 | 0.104 ± 0.096 | 0.096 ± 0.103 |
| | 05-17-10 | None | 0.000 ± 0.101 | 0.062 ± 0.117 | 0.058 ± 0.133 | 0.091 ± 0.127 | 0.119 ± 0.121 | 0.106 ± 0.117 |
| | | Rock | 0.000 ± 0.116 | 0.070 ± 0.141 | 0.423 ± 0.217 | 0.076 ± 0.131 | 0.150 ± 0.144 | 0.439 ± 0.151 |
| | | Flap | 0.019 ± 0.114 | 0.000 ± 0.095 | 0.081 ± 0.096 | 0.138 ± 0.137 | 0.110 ± 0.131 | 0.164 ± 0.102 |
| | | Flap-Rock | 0.054 ± 0.071 | 0.291 ± 0.143 | 0.323 ± 0.172 | 0.073 ± 0.031 | 0.325 ± 0.075 | 0.225 ± 0.046 |
| | 05-25-11 | None | 0.035 ± 0.100 | 0.000 ± 0.072 | 0.035 ± 0.109 | 0.113 ± 0.118 | 0.115 ± 0.118 | 0.119 ± 0.105 |
| | | Rock | 0.085 ± 0.095 | 0.067 ± 0.093 | 0.000 ± 0.119 | 0.082 ± 0.079 | 0.093 ± 0.076 | 0.115 ± 0.135 |
| 6 | 01-15-08 | None | 0.024 ± 0.106 | 0.022 ± 0.100 | 0.026 ± 0.113 | 0.116 ± 0.117 | 0.102 ± 0.124 | 0.109 ± 0.100 |
| | | Rock | 0.061 ± 0.161 | 0.070 ± 0.175 | 0.422 ± 0.241 | 0.162 ± 0.167 | 0.181 ± 0.160 | 0.408 ± 0.203 |
| | | Flap-Rock | 0.000 ± 0.108 | 0.000 ± 0.149 | 0.498 ± 0.247 | 0.079 ± 0.160 | 0.133 ± 0.194 | 0.508 ± 0.203 |
| | 01-23-08 | None | 0.054 ± 0.111 | 0.044 ± 0.103 | 0.040 ± 0.112 | 0.105 ± 0.106 | 0.105 ± 0.106 | 0.098 ± 0.103 |
| | | Rock | 0.118 ± 0.200 | 0.092 ± 0.180 | 0.480 ± 0.258 | 0.194 ± 0.156 | 0.177 ± 0.170 | 0.464 ± 0.205 |
| | 03-12-10 | None | 0.002 ± 0.096 | 0.000 ± 0.079 | 0.061 ± 0.135 | 0.088 ± 0.108 | 0.065 ± 0.105 | 0.083 ± 0.105 |
| | | Rock | 0.000 ± 0.133 | 0.000 ± 0.110 | 0.608 ± 0.220 | 0.065 ± 0.159 | 0.041 ± 0.137 | 0.456 ± 0.207 |
| | | Flap-Rock | 0.051 ± 0.093 | 0.000 ± 0.087 | 0.025 ± 0.169 | 0.203 ± 0.163 | 0.089 ± 0.093 | 0.426 ± 0.217 |

**Supplementary Table 7** — Accelerometer Median ± SD $PS_1$ Scores per Sensor, Stratified by Child, Session, and Annotation Part I

| Participant | Session | Annotation | Torso | LWrist | RWrist |
|---|---|---|---|---|---|
| 1 | 01-18-08 | None | $0.000 \pm 0.088$ | $0.000 \pm 0.042$ | $0.000 \pm 0.058$ |
| | | Rock | $0.265 \pm 0.314$ | $0.000 \pm 0.107$ | $0.000 \pm 0.129$ |
| | | Flap | $0.019 \pm 0.027$ | $0.022 \pm 0.011$ | $0.125 \pm 0.157$ |
| | | Flap-Rock | $0.324 \pm 0.178$ | $0.378 \pm 0.179$ | $0.413 \pm 0.193$ |
| | 01-25-08 | None | $0.000 \pm 0.035$ | $0.000 \pm 0.031$ | $0.000 \pm 0.058$ |
| | | Rock | $0.380 \pm 0.287$ | $0.014 \pm 0.209$ | $0.009 \pm 0.237$ |
| | | Flap | $0.017 \pm 0.070$ | $0.017 \pm 0.167$ | $0.574 \pm 0.197$ |
| | | Flap-Rock | $0.115 \pm 0.196$ | $0.124 \pm 0.173$ | $0.134 \pm 0.140$ |
| | 05-25-10 | None | $0.000 \pm 0.089$ | $0.000 \pm 0.101$ | $0.000 \pm 0.060$ |
| | | Rock | $0.642 \pm 0.286$ | $0.605 \pm 0.283$ | $0.000 \pm 0.231$ |
| | | Flap-Rock | $0.459 \pm 0.200$ | $0.590 \pm 0.186$ | $0.589 \pm 0.170$ |
| | 05-28-10 | None | $0.000 \pm 0.090$ | $0.000 \pm 0.101$ | $0.000 \pm 0.061$ |
| | | Rock | $0.633 \pm 0.288$ | $0.603 \pm 0.284$ | $0.000 \pm 0.232$ |
| | | Flap-Rock | $0.462 \pm 0.199$ | $0.589 \pm 0.184$ | $0.591 \pm 0.171$ |
| | 06-01-10 | None | $0.000 \pm 0.048$ | $0.000 \pm 0.051$ | $0.000 \pm 0.068$ |
| | | Rock | $0.474 \pm 0.320$ | $0.510 \pm 0.271$ | $0.000 \pm 0.169$ |
| | | Flap | $0.080 \pm 0.131$ | $0.042 \pm 0.104$ | $0.164 \pm 0.179$ |
| | | Flap-Rock | $0.718 \pm 0.296$ | $0.617 \pm 0.278$ | $0.175 \pm 0.193$ |
| 2 | 01-18-08 | None | $0.003 \pm 0.017$ | $0.000 \pm 0.023$ | $0.000 \pm 0.017$ |
| | | Flap | $0.013 \pm 0.033$ | $0.078 \pm 0.088$ | $0.031 \pm 0.061$ |
| | 01-24-08 | None | $0.005 \pm 0.026$ | $0.000 \pm 0.026$ | $0.003 \pm 0.049$ |
| | | Rock | $0.067 \pm 0.240$ | $0.050 \pm 0.129$ | $0.016 \pm 0.127$ |
| | | Flap | $0.050 \pm 0.045$ | $0.047 \pm 0.090$ | $0.126 \pm 0.083$ |
| | 06-04-10 | None | $0.000 \pm 0.053$ | $0.000 \pm 0.044$ | $0.000 \pm 0.089$ |
| | | Rock | $0.411 \pm 0.244$ | $0.095 \pm 0.246$ | $0.104 \pm 0.189$ |
| | | Flap | $0.000 \pm 0.003$ | $0.000 \pm 0.012$ | $0.008 \pm 0.008$ |
| | | Flap-Rock | $0.490 \pm 0.225$ | $0.000 \pm 0.187$ | $0.084 \pm 0.169$ |
| | 06-02-11 | None | $0.010 \pm 0.101$ | $0.004 \pm 0.088$ | $0.004 \pm 0.074$ |
| | | Rock | $0.000 \pm 0.086$ | $0.000 \pm 0.064$ | $0.000 \pm 0.059$ |
| 3 | 01-18-08 | None | $0.000 \pm 0.021$ | $0.000 \pm 0.035$ | $0.000 \pm 0.020$ |
| | | Flap | $0.000 \pm 0.051$ | $0.012 \pm 0.029$ | $0.015 \pm 0.181$ |
| | | Flap-Rock | $0.356 \pm 0.177$ | $0.234 \pm 0.164$ | $0.343 \pm 0.147$ |
| | 02-08-08 | None | $0.000 \pm 0.022$ | $0.000 \pm 0.025$ | $0.000 \pm 0.016$ |
| | | Flap | $0.019 \pm 0.056$ | $0.024 \pm 0.047$ | $0.000 \pm 0.062$ |
| | | Flap-Rock | $0.537 \pm 0.181$ | $0.237 \pm 0.165$ | $0.400 \pm 0.159$ |

**Supplementary Table 8** — Accelerometer Median ± SD $PS_1$ Scores per Sensor, Stratified by Child, Session, and Annotation Part II

| Participant | Session | Annotation | Torso | LWrist | RWrist |
|---|---|---|---|---|---|
| 4 | 01-17-08 | None | $0.000 \pm 0.050$ | $0.000 \pm 0.031$ | $0.000 \pm 0.048$ |
| | | Rock | $0.619 \pm 0.196$ | $0.000 \pm 0.115$ | $0.000 \pm 0.086$ |
| | | Flap | $0.093 \pm 0.084$ | $0.016 \pm 0.049$ | $0.014 \pm 0.047$ |
| | 02-07-08 | None | $0.000 \pm 0.051$ | $0.000 \pm 0.029$ | $0.002 \pm 0.045$ |
| | | Rock | $0.585 \pm 0.220$ | $0.000 \pm 0.159$ | $0.000 \pm 0.116$ |
| | | Flap | $0.058 \pm 0.072$ | $0.013 \pm 0.036$ | $0.011 \pm 0.041$ |
| | 04-27-10 | None | $0.000 \pm 0.036$ | $0.000 \pm 0.026$ | $0.000 \pm 0.059$ |
| | | Rock | $0.690 \pm 0.221$ | $0.000 \pm 0.239$ | $0.000 \pm 0.089$ |
| | | Flap | $0.122 \pm 0.055$ | $0.078 \pm 0.090$ | $0.095 \pm 0.092$ |
| | 05-11-10 | None | $0.000 \pm 0.067$ | $0.000 \pm 0.041$ | $0.000 \pm 0.050$ |
| | | Rock | $0.641 \pm 0.255$ | $0.000 \pm 0.190$ | $0.000 \pm 0.115$ |
| | | Flap | $0.081 \pm 0.067$ | $0.037 \pm 0.100$ | $0.083 \pm 0.123$ |
| 5 | 01-16-08 | None | $0.000 \pm 0.073$ | $0.000 \pm 0.046$ | $0.000 \pm 0.090$ |
| | | Rock | $0.014 \pm 0.238$ | $0.000 \pm 0.170$ | $0.001 \pm 0.193$ |
| | | Flap | $0.000 \pm 0.103$ | $0.045 \pm 0.070$ | $0.037 \pm 0.063$ |
| | | Flap-Rock | $0.298 \pm 0.177$ | $0.049 \pm 0.051$ | $0.047 \pm 0.064$ |
| | 02-08-08 | None | $0.002 \pm 0.016$ | $0.003 \pm 0.028$ | $0.003 \pm 0.020$ |
| | | Flap | $0.016 \pm 0.040$ | $0.106 \pm 0.119$ | $0.013 \pm 0.069$ |
| | 05-17-10 | None | $0.006 \pm 0.102$ | $0.020 \pm 0.082$ | $0.009 \pm 0.108$ |
| | | Rock | $0.226 \pm 0.341$ | $0.000 \pm 0.189$ | $0.000 \pm 0.178$ |
| | | Flap | $0.030 \pm 0.126$ | $0.021 \pm 0.085$ | $0.020 \pm 0.120$ |
| | | Flap-Rock | $0.467 \pm 0.242$ | $0.359 \pm 0.300$ | $0.187 \pm 0.058$ |
| | 05-25-11 | None | $0.000 \pm 0.020$ | $0.003 \pm 0.016$ | $0.000 \pm 0.022$ |
| | | Rock | $0.000 \pm 0.077$ | $0.000 \pm 0.000$ | $0.000 \pm 0.013$ |
| 6 | 01-15-08 | None | $0.000 \pm 0.015$ | $0.000 \pm 0.022$ | $0.000 \pm 0.016$ |
| | | Rock | $0.499 \pm 0.249$ | $0.037 \pm 0.181$ | $0.000 \pm 0.174$ |
| | | Flap-Rock | $0.563 \pm 0.209$ | $0.113 \pm 0.146$ | $0.075 \pm 0.125$ |
| | 01-23-08 | None | $0.000 \pm 0.011$ | $0.000 \pm 0.016$ | $0.000 \pm 0.014$ |
| | | Rock | $0.537 \pm 0.278$ | $0.000 \pm 0.169$ | $0.000 \pm 0.138$ |
| | 03-12-10 | None | $0.005 \pm 0.126$ | $0.002 \pm 0.094$ | $0.000 \pm 0.054$ |
| | | Rock | $0.461 \pm 0.224$ | $0.290 \pm 0.208$ | $0.279 \pm 0.215$ |
| | | Flap-Rock | $0.295 \pm 0.207$ | $0.279 \pm 0.174$ | $0.118 \pm 0.112$ |

**Estimated Period Summary Across Participants, Sessions, and Annotations**

**Supplementary Table 9** — Pose Landmarks Median[IQR] Period Estimation per Sensor, Stratified by Child, Session, and Annotation
Part I

| Participant | Session | Annotation | L Wrist | R Wrist | Chest | L Wrist Accel | R Wrist Accel | Chest Accel |
|---|---|---|---|---|---|---|---|---|
| 1 | 01-18-08 | None | 4.000[2.000–4.000] | 4.000[2.000–4.000] | 4.000[2.000–4.000] | 1.000[0.800–1.333] | 1.333[1.000–2.000] | 1.000[0.800–1.333] |
| | | Rock | 2.000[2.000–4.000] | 4.000[2.000–4.000] | 2.000[2.000–4.000] | 1.333[0.950–2.000] | 1.333[1.000–2.000] | 1.000[0.800–1.333] |
| | | Flap | 4.000[4.000–4.000] | 4.000[2.500–4.000] | 4.000[2.500–4.000] | 0.733[0.667–0.800] | 0.800[0.800–0.800] | 0.900[0.800–1.750] |
| | | Flap-Rock | 2.000[2.000–4.000] | 4.000[2.000–4.000] | 2.000[2.000–4.000] | 0.800[0.667–1.333] | 1.000[0.800–2.000] | 0.667[0.500–0.800] |
| | 01-25-08 | None | 4.000[2.000–4.000] | 4.000[2.000–4.000] | 4.000[2.000–4.000] | 0.800[0.571–1.000] | 1.000[0.800–1.333] | 0.800[0.571–1.000] |
| | | Rock | 2.000[1.333–4.000] | 2.000[1.333–4.000] | 2.000[1.333–4.000] | 1.333[0.800–1.333] | 1.333[1.000–1.333] | 1.333[0.800–1.333] |
| | | Flap | 2.000[1.333–4.000] | 4.000[2.000–4.000] | 2.000[0.667–4.000] | 0.571[0.571–0.667] | 2.000[2.000–4.000] | 0.667[0.571–0.667] |
| | | Flap-Rock | 2.000[1.333–4.000] | 4.000[2.000–4.000] | 4.000[2.000–4.000] | 1.000[0.800–1.333] | 1.000[0.667–1.333] | 0.667[0.500–1.000] |
| | 05-25-10 | None | 4.000[2.000–4.000] | 4.000[2.000–4.000] | 4.000[2.000–4.000] | 1.000[0.667–1.333] | 0.800[0.667–1.333] | 0.800[0.571–1.333] |
| | | Rock | 2.000[1.000–4.000] | 2.000[1.000–4.000] | 1.000[1.000–1.333] | 1.000[1.000–1.000] | 1.000[0.571–1.000] | 1.000[1.000–1.000] |
| | | Flap-Rock | 2.000[2.000–4.000] | 2.000[2.000–4.000] | 2.000[2.000–4.000] | 0.571[0.400–1.000] | 0.571[0.444–1.333] | 0.500[0.500–0.667] |
| | 05-28-10 | None | 4.000[2.000–4.000] | 4.000[2.000–4.000] | 4.000[2.000–4.000] | 1.000[0.733–1.333] | 1.333[0.800–2.000] | 0.800[0.571–1.333] |
| | | Rock | 4.000[1.333–4.000] | 2.000[2.000–4.000] | 1.000[1.000–2.000] | 1.000[1.000–1.000] | 1.333[1.000–2.000] | 1.000[1.000–1.000] |
| | | Flap-Rock | 2.000[2.000–2.000] | 2.000[2.000–4.000] | 1.000[1.000–1.833] | 1.333[1.000–2.000] | 2.000[1.000–2.000] | 1.000[0.800–1.000] |
| | 06-01-10 | None | 4.000[2.000–4.000] | 4.000[2.000–4.000] | 4.000[2.000–4.000] | 1.000[0.667–1.333] | 1.000[0.800–1.333] | 0.800[0.667–1.333] |
| | | Rock | 2.000[1.000–4.000] | 4.000[2.000–4.000] | 1.000[1.000–2.000] | 1.000[1.000–1.000] | 1.333[1.000–2.000] | 1.000[1.000–1.000] |
| | | Flap | 4.000[2.000–4.000] | 4.000[2.000–4.000] | 4.000[2.000–4.000] | 0.800[0.667–1.000] | 0.800[0.571–1.000] | 0.800[0.571–1.000] |
| | | Flap-Rock | 2.000[1.000–4.000] | 2.000[1.000–4.000] | 1.000[0.800–1.000] | 1.000[0.800–1.000] | 1.000[0.800–1.000] | 1.000[0.800–1.000] |
| 2 | 01-18-08 | None | 4.000[2.000–4.000] | 4.000[2.000–4.000] | 4.000[2.000–4.000] | 1.333[1.000–1.333] | 1.000[0.800–1.333] | 1.000[0.800–1.333] |
| | | Flap | 2.000[2.000–4.000] | 4.000[2.000–4.000] | 4.000[2.000–4.000] | 1.333[1.000–2.000] | 1.333[1.333–2.000] | 1.000[0.667–1.167] |
| | 01-24-08 | None | 4.000[2.000–4.000] | 4.000[2.000–4.000] | 4.000[2.000–4.000] | 1.333[0.800–1.333] | 0.800[0.667–1.333] | 0.800[0.667–1.333] |
| | | Rock | 2.000[2.000–4.000] | 4.000[4.000–4.000] | 2.000[2.000–2.000] | 1.000[0.800–1.333] | 4.000[4.000–4.000] | 2.000[1.833–2.000] |
| | | Flap | 2.000[1.333–4.000] | 4.000[2.000–4.000] | 2.000[2.000–4.000] | 1.333[1.000–2.000] | 1.000[0.800–1.333] | 0.667[0.571–0.800] |
| | 06-04-10 | None | 4.000[2.000–4.000] | 4.000[2.000–4.000] | 4.000[2.000–4.000] | 1.000[0.667–2.000] | 1.000[0.667–2.000] | 0.800[0.571–1.333] |
| | | Rock | 2.000[1.333–4.000] | 4.000[2.000–4.000] | 1.333[1.333–4.000] | 1.333[1.333–2.000] | 1.333[0.800–2.000] | 1.333[1.333–2.000] |
| | | Flap | 2.000[2.000–4.000] | 2.000[2.000–2.000] | 2.000[2.000–2.000] | 1.333[1.000–1.333] | 2.000[2.000–2.000] | 0.444[0.400–0.686] |
| | | Flap-Rock | 4.000[2.000–4.000] | 4.000[2.000–4.000] | 4.000[3.500–4.000] | 2.000[2.000–2.000] | 2.000[2.000–2.000] | 2.000[1.833–2.000] |
| | 06-02-11 | None | 4.000[4.000–4.000] | 4.000[4.000–4.000] | 4.000[2.000–4.000] | 1.333[0.800–2.000] | 1.333[0.667–2.000] | 1.000[0.800–1.333] |
| | | Rock | 4.000[2.000–4.000] | 4.000[4.000–4.000] | 4.000[2.000–4.000] | 1.333[0.800–2.000] | 1.333[0.800–2.000] | 2.000[1.000–2.000] |
| 3 | 01-18-08 | None | 4.000[4.000–4.000] | 4.000[2.000–4.000] | 4.000[2.000–4.000] | 1.000[0.800–1.333] | 1.000[0.800–1.333] | 1.000[0.800–1.333] |
| | | Flap | 4.000[2.000–4.000] | 4.000[2.000–4.000] | 4.000[3.500–4.000] | 1.000[0.800–1.333] | 1.000[0.800–1.083] | 1.000[1.000–1.333] |
| | | Flap-Rock | 2.000[2.000–4.000] | 4.000[2.000–4.000] | 4.000[2.000–4.000] | 0.667[0.667–1.333] | 1.000[0.667–1.333] | 0.667[0.571–1.000] |
| | 02-08-08 | None | 4.000[2.000–4.000] | 4.000[2.000–4.000] | 4.000[2.000–4.000] | 1.000[0.667–1.333] | 1.000[0.800–1.333] | 0.800[0.667–1.333] |
| | | Flap | 4.000[4.000–4.000] | 4.000[2.000–4.000] | 4.000[4.000–4.000] | 1.000[0.800–1.000] | 1.000[1.000–1.333] | 0.733[0.667–1.250] |
| | | Flap-Rock | 4.000[2.000–4.000] | 4.000[2.000–4.000] | 4.000[2.000–4.000] | 0.800[0.667–1.333] | 1.333[0.667–1.333] | 0.571[0.500–0.667] |

**Supplementary Table 10** — Pose Landmarks Median[IQR] Period Estimation per Sensor, Stratified by Child, Session, and Annotation Part II

| Participant | Session | Annotation | L Wrist | R Wrist | Chest | L Wrist Accel | R Wrist Accel | Chest Accel |
|---|---|---|---|---|---|---|---|---|
| 4 | 01-17-08 | None | 4.000[2.000–4.000] | 4.000[2.000–4.000] | 4.000[2.000–4.000] | 1.000[0.667–1.333] | 0.800[0.667–1.333] | 0.800[0.667–1.000] |
| | | Rock | 1.333[1.333–4.000] | 1.333[1.333–4.000] | 1.333[1.333–2.000] | 1.333[0.800–1.333] | 1.333[1.000–1.333] | 0.667[0.667–1.333] |
| | | Flap | 4.000[2.000–4.000] | 4.000[2.000–4.000] | 4.000[2.000–4.000] | 1.000[0.667–1.000] | 1.000[0.800–1.000] | 0.800[0.667–1.000] |
| | 02-07-08 | None | 4.000[2.000–4.000] | 4.000[2.000–4.000] | 4.000[2.000–4.000] | 1.333[0.800–2.000] | 1.000[0.667–1.333] | 0.800[0.667–1.333] |
| | | Rock | 2.000[1.333–4.000] | 2.000[1.333–2.000] | 1.333[1.333–2.000] | 0.800[0.667–1.333] | 1.333[0.800–1.333] | 0.800[0.667–1.333] |
| | | Flap | 4.000[2.000–4.000] | 4.000[2.000–4.000] | 2.000[2.000–4.000] | 1.000[0.667–1.333] | 1.000[0.800–1.333] | 0.800[0.571–1.000] |
| | 04-27-10 | None | 4.000[2.000–4.000] | 4.000[2.000–4.000] | 4.000[2.000–4.000] | 0.667[0.500–1.000] | 0.800[0.571–1.000] | 0.800[0.571–1.000] |
| | | Rock | 1.333[1.333–2.000] | 1.333[1.333–1.333] | 1.333[1.333–1.333] | 1.000[0.667–1.333] | 1.333[1.333–1.333] | 1.333[0.667–1.333] |
| | | Flap | 4.000[1.333–4.000] | 2.000[2.000–4.000] | 4.000[4.000–4.000] | 0.800[0.667–1.333] | 0.667[0.571–0.667] | 0.500[0.444–0.667] |
| | 05-11-10 | None | 4.000[2.000–4.000] | 4.000[2.000–4.000] | 4.000[2.000–4.000] | 0.800[0.667–1.333] | 1.000[0.667–1.333] | 0.800[0.571–1.333] |
| | | Rock | 2.000[1.333–2.000] | 1.333[1.333–2.000] | 1.333[1.333–1.333] | 0.800[0.667–1.000] | 1.000[0.667–1.333] | 0.800[0.667–1.333] |
| | | Flap | 2.000[2.000–4.000] | 4.000[2.000–4.000] | 4.000[2.000–4.000] | 0.667[0.571–1.000] | 1.000[0.800–1.333] | 0.667[0.571–0.800] |
| 5 | 01-16-08 | None | 4.000[2.000–4.000] | 4.000[2.000–4.000] | 4.000[2.000–4.000] | 0.800[0.667–1.333] | 0.800[0.667–1.333] | 1.000[0.800–1.333] |
| | | Rock | 2.000[2.000–4.000] | 4.000[2.000–4.000] | 2.000[2.000–4.000] | 1.333[1.000–1.333] | 1.333[1.000–2.000] | 2.000[1.333–2.000] |
| | | Flap | 4.000[2.000–4.000] | 2.000[2.000–4.000] | 4.000[2.000–4.000] | 1.333[1.000–1.333] | 1.333[1.000–1.333] | 1.333[1.333–2.000] |
| | | Flap-Rock | 4.000[2.000–4.000] | 4.000[2.000–4.000] | 2.000[2.000–2.000] | 2.000[1.333–2.000] | 2.000[2.000–2.000] | 1.333[1.333–2.000] |
| | 02-08-08 | None | 4.000[2.000–4.000] | 4.000[2.000–4.000] | 4.000[2.000–4.000] | 0.800[0.667–1.333] | 0.800[0.571–1.000] | 0.800[0.571–1.000] |
| | | Flap | 4.000[2.000–4.000] | 4.000[4.000–4.000] | 4.000[2.000–4.000] | 1.000[0.800–1.333] | 0.667[0.500–1.000] | 0.667[0.571–1.000] |
| | 05-17-10 | None | 2.000[2.000–4.000] | 4.000[2.000–4.000] | 4.000[2.000–4.000] | 1.000[0.667–1.333] | 1.333[0.800–2.000] | 0.667[0.500–1.000] |
| | | Rock | 2.000[2.000–4.000] | 4.000[2.000–4.000] | 1.333[1.333–1.333] | 1.333[1.000–1.333] | 1.333[1.000–2.000] | 1.333[1.000–1.333] |
| | | Flap | 4.000[2.000–4.000] | 4.000[2.000–4.000] | 2.000[2.000–4.000] | 1.333[0.595–2.000] | 1.000[0.667–1.333] | 0.800[0.571–1.333] |
| | | Flap-Rock | 1.333[1.333–2.000] | 1.333[1.333–4.000] | 2.000[1.333–2.000] | 1.333[1.000–1.333] | 1.333[1.333–1.333] | 1.333[1.333–1.333] |
| | 05-25-11 | None | 4.000[4.000–4.000] | 4.000[2.000–4.000] | 4.000[2.000–4.000] | 0.667[0.500–1.000] | 0.800[0.571–1.333] | 0.800[0.571–1.000] |
| | | Rock | 2.000[2.000–4.000] | 4.000[2.000–4.000] | 4.000[4.000–4.000] | 1.000[0.800–1.333] | 1.167[0.571–1.333] | 2.000[1.333–2.000] |
| 6 | 01-15-08 | None | 4.000[2.000–4.000] | 4.000[2.000–4.000] | 4.000[2.000–4.000] | 1.000[0.800–1.333] | 1.000[0.667–1.333] | 1.000[0.667–1.333] |
| | | Rock | 2.000[1.333–4.000] | 2.000[1.333–4.000] | 1.333[1.333–2.000] | 1.333[1.333–1.333] | 1.333[1.000–1.333] | 1.333[1.333–1.333] |
| | | Flap-Rock | 2.000[2.000–4.000] | 4.000[2.000–4.000] | 1.333[1.000–2.000] | 2.000[1.333–2.000] | 1.333[1.333–2.000] | 1.000[1.000–1.333] |
| | 01-23-08 | None | 4.000[2.000–4.000] | 4.000[2.000–4.000] | 4.000[2.000–4.000] | 0.800[0.571–1.333] | 0.800[0.571–1.000] | 1.000[0.800–1.333] |
| | | Rock | 2.000[1.333–4.000] | 2.000[1.333–4.000] | 1.333[1.333–1.333] | 1.333[1.333–1.333] | 1.333[1.333–1.333] | 1.333[1.333–1.333] |
| | 03-12-10 | None | 4.000[2.000–4.000] | 4.000[2.000–4.000] | 4.000[2.000–4.000] | 1.333[0.800–2.000] | 1.000[0.800–1.333] | 0.800[0.571–1.333] |
| | | Rock | 4.000[2.000–4.000] | 4.000[2.000–4.000] | 1.333[1.333–1.333] | 2.000[1.333–4.000] | 2.000[1.333–2.000] | 1.333[1.333–1.333] |
| | | Flap-Rock | 4.000[2.000–4.000] | 4.000[2.000–4.000] | 4.000[1.000–4.000] | 1.333[0.458–2.000] | 0.667[0.500–1.000] | 0.571[0.500–0.800] |

**Supplementary Table 11** — Accelerometer Median[IQR] Period Estimation per Sensor, Stratified by Child, Session, and Annotation Part I

| Participant | Session | Annotation | Torso | L Wrist | R Wrist |
|---|---|---|---|---|---|
| 1 | 01-18-08 | None | 3.700 [3.690–3.700] | 3.700 [3.100–3.700] | 3.690 [3.100–3.700] |
| | | Rock | 1.700 [1.298–3.690] | 3.690 [2.896–3.700] | 2.900 [1.302–3.700] |
| | | Flap | 3.700 [3.700–3.700] | 3.690 [3.690–3.690] | 1.097 [1.097–1.097] |
| | | Flap-Rock | 1.297 [1.100–1.700] | 1.297 [1.100–1.300] | 1.297 [1.100–1.300] |
| | 01-25-08 | None | 3.700 [3.690–3.700] | 3.700 [3.690–3.700] | 3.700 [3.681–3.700] |
| | | Rock | 1.696 [1.298–3.690] | 1.704 [1.300–3.700] | 2.908 [1.298–3.700] |
| | | Flap | 3.695 [3.693–3.698] | 3.700 [3.700–3.700] | 2.390 [1.745–3.035] |
| | | Flap-Rock | 1.696 [1.300–3.104] | 1.100 [1.099–1.302] | 1.103 [1.100–1.300] |
| | 05-25-10 | None | 3.700 [3.700–3.707] | 3.700 [3.100–3.710] | 3.700 [3.108–3.700] |
| | | Rock | 1.103 [1.100–1.201] | 1.100 [1.100–1.103] | 3.700 [1.201–3.705] |
| | | Flap-Rock | 1.103 [1.100–1.303] | 1.103 [1.100–1.300] | 1.100 [0.900–1.300] |
| | 05-28-10 | None | 3.700 [3.700–3.700] | 3.700 [3.100–3.710] | 3.700 [3.108–3.700] |
| | | Rock | 1.100 [1.100–1.251] | 1.100 [1.100–1.103] | 3.700 [1.152–3.700] |
| | | Flap-Rock | 1.103 [1.100–1.300] | 1.103 [1.100–1.300] | 1.100 [0.900–1.300] |
| | 06-01-10 | None | 3.700 [3.700–3.710] | 3.700 [3.100–3.700] | 3.700 [3.100–3.707] |
| | | Rock | 1.101 [1.100–1.103] | 1.101 [1.100–1.103] | 3.400 [2.651–3.700] |
| | | Flap | 3.700 [3.700–3.710] | 3.700 [2.900–3.700] | 1.700 [1.103–3.700] |
| | | Flap-Rock | 1.100 [1.100–1.700] | 1.103 [1.100–1.700] | 1.103 [0.900–1.300] |
| 2 | 01-18-08 | None | 3.700 [3.690–3.700] | 3.690 [3.092–3.700] | 3.700 [3.690–3.700] |
| | | Flap | 3.700 [3.700–3.700] | 1.297 [1.097–1.300] | 3.690 [1.300–3.710] |
| | 01-24-08 | None | 3.700 [3.690–3.700] | 3.100 [2.894–3.700] | 3.695 [3.690–3.700] |
| | | Rock | 1.803 [1.552–2.055] | 1.797 [1.548–2.045] | 1.795 [1.546–2.045] |
| | | Flap | 3.700 [3.690–3.702] | 1.200 [1.099–1.403] | 1.200 [1.000–2.000] |
| | 06-04-10 | None | 3.700 [3.100–3.700] | 3.108 [2.200–3.700] | 3.004 [2.300–3.700] |
| | | Rock | 1.302 [1.251–1.700] | 1.700 [1.303–2.451] | 1.802 [1.300–2.956] |
| | | Flap | 3.700 [3.700–3.700] | 3.705 [3.702–3.707] | 2.000 [1.850–2.150] |
| | | Flap-Rock | 1.700 [1.700–1.700] | 2.700 [2.200–3.200] | 1.500 [1.400–1.600] |
| | 06-02-11 | None | 2.908 [1.700–3.700] | 2.306 [1.303–3.108] | 2.303 [1.300–3.698] |
| | | Rock | 2.306 [2.300–2.900] | 2.306 [2.300–2.908] | 2.306 [1.992–2.906] |
| 3 | 01-18-08 | None | 3.700 [3.690–3.700] | 3.690 [3.614–3.700] | 3.700 [3.690–3.700] |
| | | Flap | 3.700 [3.690–3.700] | 3.100 [1.097–3.690] | 1.300 [1.201–3.399] |
| | | Flap-Rock | 1.198 [1.098–1.303] | 1.198 [1.100–1.299] | 1.200 [1.100–1.297] |
| | 02-08-08 | None | 3.700 [3.690–3.700] | 3.700 [3.690–3.700] | 3.700 [3.690–3.700] |
| | | Flap | 3.700 [3.690–3.700] | 3.092 [3.092–3.096] | 3.100 [3.000–3.695] |
| | | Flap-Rock | 1.200 [1.100–1.297] | 1.100 [1.100–1.297] | 1.300 [1.100–1.401] |

**Supplementary Table 12** — Accelerometer Median[IQR] Period Estimation per Sensor, Stratified by Child, Session, and Annotation Part II

| Participant | Session | Annotation | Torso | L Wrist | R Wrist |
|---|---|---|---|---|---|
| 4 | 01-17-08 | None | 3.700 [3.690–3.700] | 3.690 [3.635–3.700] | 3.700 [3.690–3.700] |
| | | Rock | 1.300 [1.300–1.300] | 3.700 [3.681–3.700] | 3.700 [3.690–3.700] |
| | | Flap | 3.700 [3.690–3.700] | 2.300 [1.300–2.908] | 2.294 [1.300–2.900] |
| | 02-07-08 | None | 3.690 [3.102–3.700] | 3.690 [2.944–3.700] | 3.700 [3.100–3.700] |
| | | Rock | 1.300 [1.297–1.696] | 3.690 [3.092–3.700] | 3.700 [3.102–3.700] |
| | | Flap | 3.700 [3.690–3.700] | 1.700 [1.293–2.900] | 2.093 [1.297–2.898] |
| | 04-27-10 | None | 3.700 [3.700–3.710] | 3.108 [2.271–3.700] | 3.700 [2.904–3.700] |
| | | Rock | 1.300 [1.300–1.303] | 1.504 [1.303–3.700] | 3.700 [3.100–3.710] |
| | | Flap | 3.100 [3.100–3.104] | 2.900 [2.600–2.900] | 2.300 [1.701–2.600] |
| | 05-11-10 | None | 3.700 [3.700–3.710] | 3.700 [3.108–3.710] | 3.700 [3.100–3.700] |
| | | Rock | 1.300 [1.300–1.696] | 3.700 [1.700–3.700] | 3.700 [2.908–3.700] |
| | | Flap | 3.700 [3.700–3.700] | 1.300 [1.100–2.400] | 1.300 [1.300–2.002] |
| 5 | 01-16-08 | None | 3.700 [3.690–3.700] | 3.690 [3.647–3.700] | 3.700 [3.690–3.700] |
| | | Rock | 2.103 [1.700–2.948] | 3.381 [2.200–3.690] | 3.000 [1.850–3.683] |
| | | Flap | 3.690 [3.690–3.700] | 1.696 [1.297–1.700] | 1.303 [1.300–2.306] |
| | | Flap-Rock | 1.700 [1.694–1.704] | 1.698 [1.299–1.700] | 1.700 [1.597–1.700] |
| | 02-08-08 | None | 3.700 [3.690–3.700] | 3.700 [3.690–3.700] | 3.700 [3.690–3.700] |
| | | Flap | 3.700 [3.700–3.700] | 1.700 [1.100–3.690] | 3.700 [3.690–3.700] |
| | 05-17-10 | None | 3.700 [3.700–3.700] | 3.700 [3.700–3.710] | 3.700 [3.690–3.705] |
| | | Rock | 1.303 [1.103–3.700] | 3.700 [3.100–3.710] | 3.700 [2.885–3.700] |
| | | Flap | 3.700 [3.698–3.700] | 3.104 [1.904–3.700] | 3.681 [1.702–3.710] |
| | | Flap-Rock | 1.704 [1.704–1.704] | 1.700 [1.700–1.700] | 1.700 [1.700–1.700] |
| | 05-25-11 | None | 3.705 [3.700–3.710] | 3.710 [3.700–3.710] | 3.710 [3.700–3.710] |
| | | Rock | 3.400 [2.950–3.700] | 3.700 [3.529–3.700] | 3.572 [3.041–3.695] |
| 6 | 01-15-08 | None | 3.700 [3.690–3.700] | 3.690 [3.681–3.700] | 3.700 [3.690–3.700] |
| | | Rock | 1.300 [1.103–1.303] | 2.300 [1.297–3.688] | 3.700 [2.900–3.700] |
| | | Flap-Rock | 1.103 [1.100–1.300] | 1.103 [1.097–1.300] | 1.103 [1.100–1.303] |
| | 01-23-08 | None | 3.700 [3.700–3.700] | 3.690 [3.690–3.700] | 3.700 [3.690–3.700] |
| | | Rock | 1.300 [1.297–1.303] | 3.092 [1.303–3.690] | 3.700 [3.690–3.700] |
| | 03-12-10 | None | 3.700 [3.700–3.710] | 3.700 [3.100–3.710] | 3.359 [2.908–3.700] |
| | | Rock | 1.300 [1.300–1.303] | 1.300 [1.278–1.303] | 1.300 [1.278–1.303] |
| | | Flap-Rock | 1.101 [0.800–1.251] | 1.100 [0.801–1.253] | 1.300 [1.150–1.303] |
